# Dual Role of Microglial TREM2 in Neuronal Degeneration and Regeneration After Axotomy

**DOI:** 10.1101/2025.08.06.668924

**Authors:** Tana S. Pottorf, Elizabeth L. Lane, Zoë Haley-Johnson, Desirée N. Ukmar, Veronica Amores- Sanchez, Patricia M. Correa-Torres, Francisco J. Alvarez

**Affiliations:** Department of Cell Biology, Emory University School of Medicine, Atlanta, GA 30322, USA; Department of Biology, University of Puerto Rico at Ponce, PR00931, Puerto Rico

**Keywords:** Peripheral Nerve Injury, Motoneuron, Microglia, TREM2, CD68, Microglia morphology, Phagocytosis

## Abstract

Ventral horn microglia in the spinal cord proliferate after nerve injuries and migrate towards the cell bodies of injured motoneurons surrounding them. However, the significance of microglia enwrapping axotomized motoneurons has remained unclear. Moreover, some injured motoneurons degenerate while others regenerate. In mice spinal cords we found that each motoneuron fate associates with microglia of different activation profiles. Microglia surrounding degenerating motoneurons form cell clusters that fully envelop the cell body and express high TREM2 and large CD68 granules, with female microglia expressing higher levels. Microglia surrounding motoneurons undergoing regeneration remain individualized and also upregulate TREM2 and CD68, but to a lesser extent than microglia around degenerating motoneurons. Removal of TREM2, either globally throughout development or specifically in microglia prior to nerve injuries, reduces p-SYK signaling and CD68 expression in all activated microglia, but more so inside microglia forming tight cell clusters around degenerating motoneurons. This effect is also larger in females. TREM2 absence did not prevent microglia clustering around degenerating motoneurons but prevented the loss of some small MNs. In addition, TREM2 depletion interfered with the retrograde cell body chromatolytic reaction that is characteristic of regenerating motoneurons and delayed muscle reinnervation. We conclude that within the same motor pools, TREM2 facilitates microglia removal of some degenerating motoneurons while it facilitates regeneration of other motoneurons. The signals that direct the development of these different microglia phenotypes over degenerating and regenerating motoneurons, as well as the mechanisms that induce degeneration in some motoneurons while most others regenerate, remain to be investigated.

**Significance Statement:** Microglia frequently enwrap neurons undergoing cellular stress being one example the microglia reaction around motoneurons axotomized after nerve injuries. The significance of this microglia-neuron relationship is unclear. We found that microglia surrounding axotomized motoneurons upregulate TREM2, but with differences depending on whether microglia associated with regenerative or degenerative motoneurons. Loss-of-function experiments showed that TREM2 promotes removal of some degenerating motoneurons while facilitates the regeneration of others. We conclude that microglia TREM2 serves a dual function depending on the motoneuron health state. This knowledge is critical for designing future therapies that aim to improve motoneuron regeneration or preservation by altering TREM2 function.

## Introduction

In a landmark 1968 paper Blinzinger and Kreutzberg (Blinzinger & Kreutzberg, 1968) reported proliferating microglia surrounding facial motoneurons (MNs) after axotomy following injuries to the facial nerve. They proposed that the microglia reaction was protective working through the removal of synapses surrounding the cell bodies of injured MNs. This phenomenon was named “synaptic stripping” because microglia processes were observed interposed between synaptic boutons and the MN surface without synaptic engulfment. Now, it is well accepted that microglia can remove synapses via phagocytosis or trogocytosis during normal developmental circuit refinement and in disease (Hong et al., 2016; Pereira-Iglesias et al., 2025; Piccioni et al., 2021; Rajendran & Paolicelli, 2018; Stevens et al., 2007; Vecchiarelli et al., 2024; Weinhard et al., 2018; Wilton et al., 2019). Nevertheless, the Blinzinger and Kreutzberg original hypothesis that microglia remove synapses from the cell bodies of axotomized MNs has been repeatedly challenged (Berg et al., 2013; Kalla et al., 2001; Rotterman et al., 2019; Rotterman et al., 2024; Salvany et al., 2021; Svensson & Aldskogius, 1993a). This raises questions about the significance of microglia surrounding injured MN cell bodies.

We reported that axotomized MNs orchestrate the microglia reaction around them by upregulating Colony Stimulating Factor 1 (CSF1) following peripheral nerve injury (PNI) (Rotterman et al., 2019; Rotterman et al., 2024). Injury severity extends the time-course of CSF1 expression in MNs prolonging microglia activation and promoting microglia to switch towards proinflammatory phenotypes that express C-C chemokine ligand 2 (CCL2). CCL2-expressing microglia are associated with significant sensorimotor circuit plasticity (Rotterman et al., 2024) that results in long-lasting motor deficits (Alvarez et al., 2010; Alvarez et al., 2020). Thus, ventral horn microglia can display a variety of activated microglia phenotypes dynamically regulated by injury severity and time after injury. After nerve transections requiring surgical repair two distinct microglia-MN interactions were reported (Rotterman & Alvarez, 2020). More commonly, microglia migrate towards the cell bodies of MNs that are in the process of regenerating and surround them while maintaining cell individuality. A second, less frequent, phenotype consists of tight cell clusters of highly phagocytic ameboid microglia with retracted processes, which completely surround degenerating MNs and attract cytotoxic T-cells (CD8+) (Rotterman & Alvarez, 2020). The significance and mechanisms regulating these two types of microglia-MN interactions and therefore, MN fates after injury, regeneration or degeneration, are unknown.

We focused on Triggering Receptor Expressed on Myeloid Cells 2 (TREM2) because of its function promoting neuronal phagocytosis by microglia (Takahashi et al., 2005) and because expression of a protein essential for TREM2 function, DNAX Activating Protein of 12 kDa (DAP12), is regulated by CSF1 after PNI (Guan et al., 2016). We hypothesized that TREM2, which recognizes a variety of “eat me” signals (Rueda-Carrasco et al., 2023; Scott-Hewitt et al., 2020; Shirotani et al., 2019) may principally switch the properties of microglia associated with degenerating MNs towards phagocytic phenotypes. However, microglia TREM2/DAP12 mechanisms have other roles beyond phagocytosis in both healthy and diseased states, including synapse deletion (Filipello et al., 2018; Jay et al., 2019; Matteoli, 2024; Rueda-Carrasco et al., 2023; St-Pierre et al., 2020; Vecchiarelli et al., 2024; Zhong et al., 2023) and metabolic support of microglia and neurons (Tagliatti et al., 2024). These functions are critical for regeneration or degeneration of MNs after nerve injuries. We therefore set out to analyze TREM2 expression in microglia surrounding injured motoneurons and determine TREM2 functions using two knockout models: a global knockout and a conditional knockout in which *trem2* was selectively deleted in microglia before the nerve injury. Our findings revealed a nuanced and complex role for TREM2 with different functions that depend on the health state of the MN microglia interact with. In some cases, TREM2 promoted the regeneration of axotomized MNs, whereas when interacting with MNs undergoing degeneration TREM2 upregulates phagocytic markers like CD68. Notably, TREM2 deletion did not significantly alter microglial proliferation, activation-related morphological changes, or migration towards injured MNs.

## Materials and Methods

### Ethical Statement

All animal procedures were approved by the Institutional Animal Care and Use Committee at Emory (Protocol #201700620) and followed the NIH Guidelines for the use of animals in research. All animal procedures, analyses, and husbandry were performed at Emory University. We followed ARRIVE guidelines during experimental design and performance.

### Experimental Animals

Mice used in this manuscript were from a C57Blk/6 J/N mixed background. The C57BL/6-Tmem119^em2Gfng^/J and C57BL/6J-TREM2^em2Adiuj^/J mice were bred to create a global TREM2 knockout (GKO) mouse with microglia-specific enhanced Green Fluorescent Protein (eGFP) labeling. The C57BL/6-Tmem119^em1(cre/ERT2)Gfng^/J, C57BL/6J-Trem2^tm1c(EUCOMM)Wtsi/Adiuj^/J, and B6.Cg-Gt (ROSA) ^26Sortm9(CAG-tdTomato)Hze^/J were bred to create a Tmem119-2A-CreERT2:: TREM2 flox/flox:: Rosa26 tdTomato mouse in which after tamoxifen induced cre recombination Tmem119-specific microglia are labeled with tdTomato and simultaneously knocked out TREM2 (conditional knockout: CKO). Finally, *B6.129P-Cx3cr1^tm1Litt^/J* mice are *CX3CR1^eGF^*^P^ knock-in knock-out mice bred in heterozygosis to label microglia with GFP. All other mice were bred in homozygosis. Genetic mouse lines and their usage in this study are summarized in **Table 1**.

We found no significant differences in the number of ventral horn microglia labeled with antibodies against Ionized calcium Binding Adaptor molecule 1 (IBA1) versus CX3CR1^eGFP^ or Tmem119^eGFP^ (**Figure 4C**). This allowed us to use different microglia genetic or immunocytochemical labels interchangeably and best adapted to the needs of each technique or experiment. Transgenic mice were housed with littermates, standard chow, and 12-hour light-dark cycles. All mice were housed in auto-water racks except for mice treated with tamoxifen which were group-housed in static racks after treatment.

### Tamoxifen Injections

To induce cre recombination, CKO mice were injected with tamoxifen (Sigma Aldrich, cat: T5648) once daily for five consecutive days starting at postnatal day 28 (intraperitoneal (i.p.) 2 mg/kg, diluted in corn oil; Ward’s Science, cat: 470200-112).

### Intramuscular Injections for Retrograde Motoneuron Labeling

For all survival surgeries, mice were anesthetized via isoflurane inhalation (4% induction and 2% maintenance) and received 0.05 mg/kg buprenorphine subcutaneously to diminish postoperative pain. The left lateral gastrocnemius (LG) muscle was exposed and targeted for injections after a 4-6 mm skin cut in adult and 2-3 mm in neonates. Four μL of 2% Fast Blue (Polyscience, cat: 17740-1) was injected using a 25G Hamilton syringe in two-month-old animals. Skin wounds were closed with Vetbond®, in both neonates and adults.

### Sciatic Cut Ligation and Sham Surgeries

Nerve surgeries were performed in adult mice of two to four months of age and both sexes. When Fast Blue intramuscular injections were used, a minimum of five days was waited before performing nerve surgeries to allow time for retrograde transport of the tracer. A skin incision was made in the medial left thigh, parallel to the femur. Then the biceps femoris was retracted by blunt dissection of the fascia to expose the sciatic nerve. The sciatic nerve was isolated and either left intact (sham surgery) or ligated with a 5-0 non-absorbable suture and cut distally (cut-ligation). The muscle was reconnected with absorbable 5-0 sutures, and the skin was closed with non-absorbable 5-0 sutures and Vetbond®. Anesthesia (isoflurane) and analgesia (buprenorphine) were as above.

### Sciatic Cut Repair With or Without Fast Blue Nerve Soak

Mice undergoing sciatic nerve cut-repair surgeries frequently received a nerve soak with Fast Blue at the time of the injury. The sciatic nerve was exposed at mid-thigh in two to four-month-old mice and then partially lifted to insert a 1 mm^2^ silastic scaffolding matrix (Dow Corning #501-1) underneath. Fibrin Glue (Akhter et al., 2019) was added to the nerve on the scaffolding mesh and allowed to partially harden. The nerve was then transected with fine surgical scissors. Fibrin Glue maintained alignment between the nerve stumps, and at this time, 1 μL of 2% Fast Blue was added to the exposed nerve stumps. Additional Fibrin Glue was added on top of the transected nerve before closing. Muscles and skin were repaired and closed as above. Anesthesia (isoflurane) and analgesia (buprenorphine) were as above.

### Transcardial Perfusion with Fixatives and Tissue Collection

Mice were anesthetized with a terminal dose of Euthasol (100 mg/kg) at 3, 7, 14, 21, or 28 days post-injury (dpi) and transcardially perfused with 5 mL of heparin pre-perfusion rinse followed by 15 mL of 4% paraformaldehyde (PFA) prepared in 0.1M phosphate buffer (pH=7.4). The spinal cords were harvested and placed in 4% PFA for post-fixation. Spinal cord tissues used for immunohistochemistry were post-fixed for four hours at 4°C, followed by a minimum of 24 hours in 30% sucrose. Spinal cord tissue used for RNAscope was post-fixed for 24 hours at 4°C, followed by 24 hours in 10% sucrose, 24 hours in 20%, and 24 hours in 30% sucrose. Muscle tissue was post-fixed for 4 hours at 4°C, followed by 24 hours in 10% sucrose, 24 hours in 20%, and 24 hours in 30% sucrose.

### Immunohistochemistry and Confocal Microscopy

Transverse sections (50-75 μm thick, depending on experiments) from the Lumbar 3 (L3) to the L6 segment were obtained on a freezing sliding microtome and collected free-floating. Free-floating sections were subjected to a one-hour block in 10% Normal Donkey Serum diluted in 0.01M Phosphate Buffer Solution with 0.03% Triton (PBST). The sections were then incubated in different mixtures of primary antibodies, all diluted in PBST. Primary antibody incubations were carried out overnight for 50 µm thick sections or 48 hours for thicker 75 µm sections, both at room temperature (different antibody combinations explained in each analysis). The following day, immunoreactive sites were revealed with species-specific secondary antibodies conjugated to different fluorochromes (2-3 hours at room temperature), and after washes in PBS, the sections were mounted on glass slides and cover-slipped with Vectashield with DAPI (Vector Labs, cat: H-1200-10), or without (Vector Labs, cat: H-1000-10). Details of antibodies used and their concentrations are listed in **Table 2**. Confocal microscopy was performed with 10X (NA 0.4), 20X (NA 0.7), and 60X (oil, NA 1.35) objectives in a Fluoview FV1000 Olympus microscope. All images were 1024X1024 pixels in size. Digital zoom and z-step sizes varied depending on experiments. Antibody combinations and imaging conditions are described in each analysis.

Ipsilateral LG muscle tissue was isolated and compressed as described before (Wood et al., 2025). The compressed LG muscles were embedded in OCT (Optimal Cutting Temperature Compound) and frozen before cutting 75 μm sections with a cryostat. Sections were air dry on the slides and kept at -20°C for a minimum of overnight before immunohistochemistry as described above. The slides containing sections were washed with PBS to remove the OCT and subjected to one hour block in 10% Normal Donkey Serum diluted in PBST followed by overnight incubation (room temperature) in a mix of anti-VAChT and anti-Neurofilament heavy chain (NFH) antibodies diluted in PBST (dilutions, host, and sources are in **Table 2**). α-Bungarotoxin (α-BTx) conjugated to Alexa 488 or 647 was added to the primary antibody mixture. The following day, immunoreactive sites were revealed with appropriate fluorochrome-conjugated species-specific antibodies incubated for two hours at room temperature. Finally, the slides were washed in PBS and cover-slipped with Vectashield with DAPI (Vector Labs, H-1200-10).

### Microglia Sholl Analysis

Tmem119^eGFP^ microglia associated with MNs retrogradely labeled with Fast Blue were imaged with confocal microscopy (60x objective, 2x digital zoom, 0.5 μm step size) in animals with sham or sciatic nerve cut and ligation surgeries (N=6 shams and 6 injured, equal sex). We divided microglia into four groups according to microglia location in relation to retrogradely labeled MNs in animals with 1) sham surgeries or the 2) contralateral spinal cord in animals with nerve injuries, and two locations in the spinal cord ipsilateral to nerve injuries, 3) associated with injured Fast Blue motoneurons that were not degenerating (“On MN” microglia) and 4) integrated in microglia cell clusters associated with degenerating motoneurons. We denominate this as death cluster for short and define these as DC microglia. From here on, “death cluster” will refer to the cluster, and “DC” will refer to the individual microglia within these clusters. In each animal, we analyzed five microglia per condition (sham, contralateral, On MN, or DC). The resulting sample sizes are shown in **Data S1** in support of **Figure 1E**. Individual microglia were semi-automatically traced and skeletonized in IMARIS by thresholding the fluorescent eGFP signal, followed by cell isolation and 3D reconstruction using the filament tracer module (n=5 microglia per animal, 30 per experimental condition). Sholl analyses for microglia processes transecting spheres increasing by 10 μm in distance from the cell body center were performed in IMARIS.

**Figure 1.**
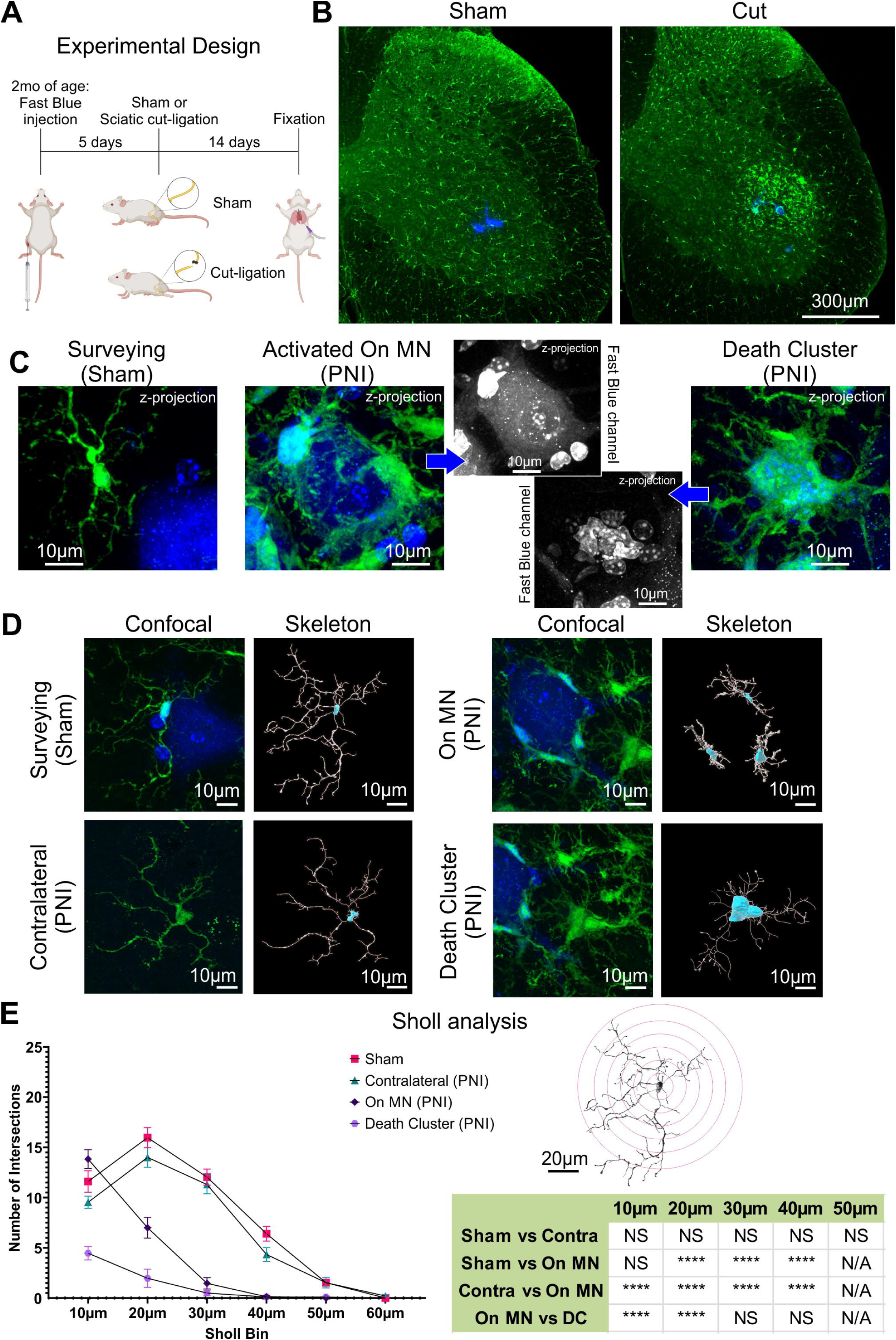
Activated microglia display distinct morphologies depending on the health state of the motoneuron they interact with after nerve injury. **A**) Experimental design. The lateral gastrocnemius muscle of Tmem119^eGFP^ mice was injected with Fast Blue (FB) five days before sciatic nerve surgeries, and the animals were analyzed 14 days post-surgery. We prepared sham and cut-ligation mice (N=6 mice each, equal sex). **B**) Representative low magnification z-stack 2D projection images of lumbar 5 (L5) spinal cord sections (50 µm thick) from sham or cut-ligation mice. eGFP microglia (green) proliferate and surround FB motoneurons (MNs) after nerve injury, but not in sham animals. **C**) High-magnification 2D projections (25 µm thick z-stacks) of microglia (green) close to FB MNs in a sham animal, or after PNI on an injured MN or in a death cluster. The Fast Blue channel in injured MNs is shown in black and white to demonstrate the morphologies of healthy, likely regenerating, vs degenerating MNs. Fast Blue is incorporated by microglia in both types of microglia-MN interactions. **D**) High magnification confocal images and cell skeletonizations of surveying microglia in shams and contralateral cut-ligation animals, and On MN and DC microglia ipsilateral to cut-ligation animals. **E**) Sholl analysis of microglia in each group showing average number of intersections in each Sholl bin ±SEM (n=30 microglia per group, 5 microglia per animal). Summary of two-way ANOVA comparisons for Sholl distance and microglia type (p<0.0001 for both) followed by Bonferroni multiple comparisons. The table shows some relevant comparisons; ***p<0.001, ***p<0.0001. Statistical details in **Data S1**.

### Analysis of TREM2 Immunofluorescence

TREM2 immunofluorescence was performed in Tmem119^eGFP^ mice with Fast Blue MNs retrogradely labeled from the LG muscle (N=8 sham and 8 with sciatic-nerve injury and ligation; equal sex). Animals were analyzed 14 days after surgery. Confocal z-stack images were taken from triple fluorescent sections with Fast Blue positive MNs, eGFP or tdTomato microglia and TREM2 immunofluorescence (Cy5) using the 60x objective, 2x digital zoom, and 1 μm step size. Both fluorescent proteins were amplified with antibodies in the same channel but note that for consistency illustrating microglia genetic labeling we sometimes pseudo-colored tdtomato as green in the figures. We imaged between 20 to 26 sham, contralateral, and On MN microglia per animal. Fewer DC microglia were sampled per animal. TREM2 protein signal-to-noise ratio was calculated by examining each optical plane through the z-stack and identifying the location of the microglia cell body mid-cross section. A Region of Interest (ROI) was drawn around the membrane using ImageJ and excluding the Golgi that is always strongly stained (Sessa et al., 2004). The average fluorescence intensity was measured in the ROI and similar-sized ROIs were sampled in a background area near each microglia. To estimate cell body fluorescence intensity over background, we use the formula sample – background)/(background. Individual microglia outlier estimates were identified via the ROUT method (Q=1%) using Prism (GraphPad). The resulting sample sizes are shown in **Data S1** in support of **Figures 2B and C**.

**Figure 2.**
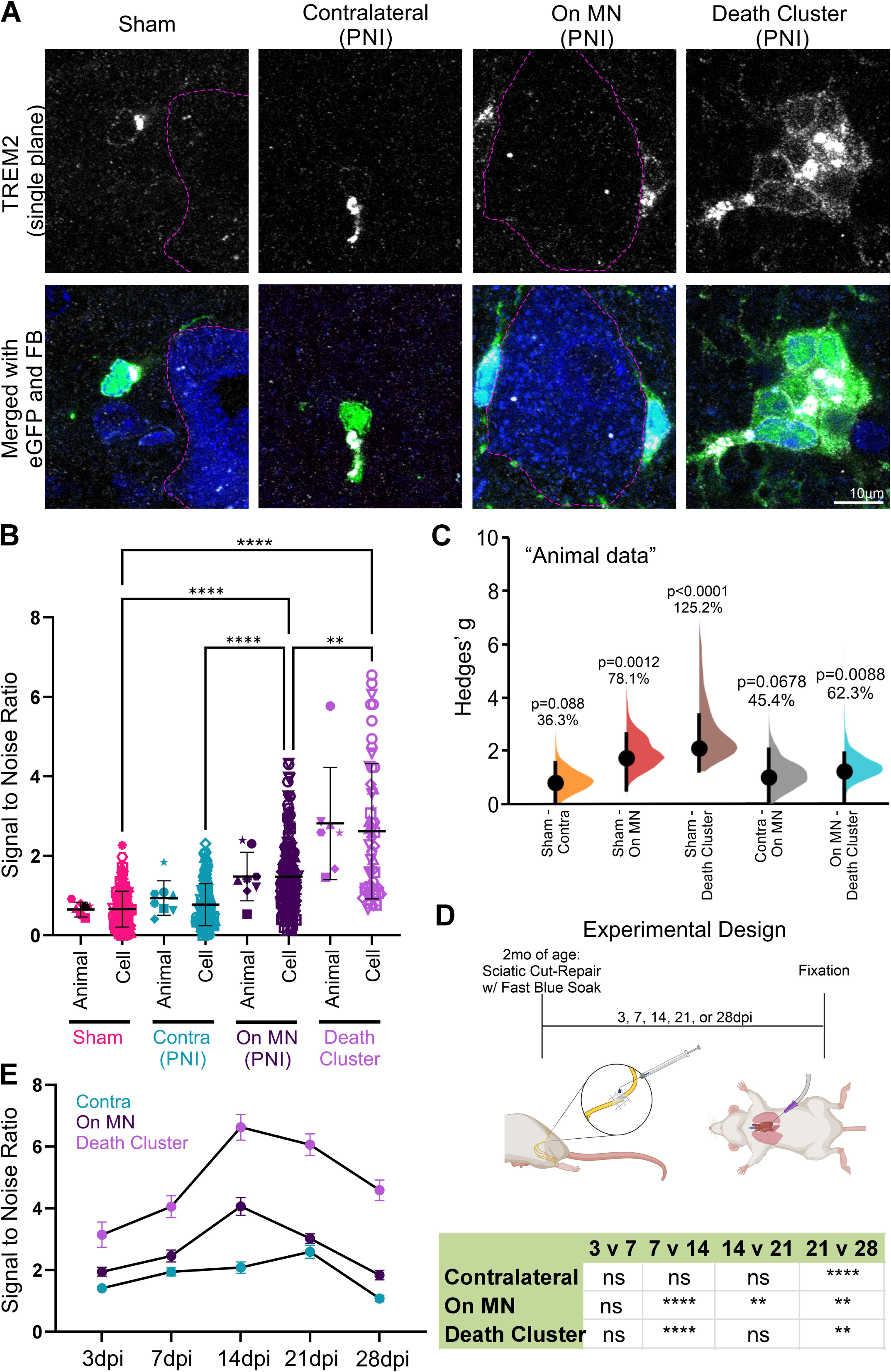
TREM2 expression is highest at 14 days post-injury and in Death Cluster microglia. **A**) Single plane confocal images of Tmem119^eGFP^ microglia (green) immunolabeled for TREM2 (white) and close to FB motoneurons (MNs). Microglia display high TREM2 in patches juxtaposed to the nucleus corresponding to the Golgi (Sessa et al., 2004). Cytoplasmic and membrane TREM2 immunofluorescence increased in activated microglia and was maximal in DC microglia. **B**) TREM2 signal calculated in the membrane and cytosol, corrected for background noise. Individual microglia (Cell) and animal (Animal) averages are shown (n=17-26 microglia per animal and condition, N=8 mice (equal sex) per condition). Each symbol represents one animal or microglia cell. Symbol shapes identify individual animals. Mean±SD is represented. For simplicity, only the most salient Dunn’s post-hoc multiple comparisons of Cell data are shown (**p<0.01, ****p<0.0001) (see **Data S1** for details and other statistical comparisons). **C**) Animal averages analyzed with estimation statistics for effect sizes estimated by Hedge’s g of mean differences. Dots show mean differences; vertical error bars 95% confidence intervals. Percentage differences and significance (permutation t-tests) are indicated on each graph. **D**) Experimental design for time course analysis of TREM2 expression after sciatic nerve cut and repair surgeries. N=4 animals at each dpi, n=20 microglia analyzed per animal for On MN and contralateral MNs. Each time point includes 72 to 79 microglia. Death cluster recovery varied: 3 dpi n=16, N=1; 7 dpi n=29, N=2; 14 dpi n=80, N=4; 21 dpi n=48 N=3; 28 dpi n=37, N=2. **E**) TREM2 immunofluorescence at different days post-injury (dpi) in microglia located in the contralateral ventral horn, or ipsilateral On MN and DC microglia. Mean±SEM of the Cell data. TREM2 expression peaks at 14 dpi in microglia ipsilateral to the injury with DC microglia exhibiting higher TREM2 that takes longer to decay. Two-way ANOVA (type of microglia and dpi) showed significance for both variables (p<0.001). The table on the right shows significant differences at different time points (Bonferroni tests, details of samples and statistics in **Data S1**).

To confirm TREM2 protein absence in GKO and CKO mice, we used the same method, although in this case microglia identification was also supported by immunolabeling with IBA1 antibodies.

### CX3CR1 *trem2* RNAscope co-detection and Analysis

RNAscope multiplex fluorescent v2 assay (ACDBio, Cat:323100) was performed in CX3CR1^eGFP^ animals and combined with eGFP immunofluorescence-integrated co-detection workflow on fixed-frozen tissue sections according to the suggested Advance Cell Diagnostics protocol (acdbio.com). Isolated tissues were fixed and cryoprotected as explained above, and then 16 μm-thick cryostat sections were obtained from L3 to L5 segments, collected on slides, air-dried, and kept at -20°C until use. The reagents used to perform RNAscope were as follows: RNAscope Intro Pack for Multiplex Fluorescent Reagent Kit v2-Mm (ACDBio, Cat: 323136), RNA-Protein Co-detection Ancillary Kit (ACDBio, Cat:323180), RNAscope probe-Mm-TREM2 (ACDBio, Cat: 404111), Opal 570 Reagent Pack (Akoya Bioscience, Cat: FP1488001KT). Genetically encoded eGFP was amplified with primary antibody, chicken anti-eGFP (Aves, 1:50 dilution, Table 2), and secondary antibody, FITC anti-chicken IgY (1:50 dilution, Table 2). Genetically encoded tdTomato was amplified with a goat anti-mCherry antibody (MyBioSource, 1:50 dilution, **Table 2**) followed by Cy3 anti-goat IgG (1:50 dilution, **Table 2**). Antibodies were used more concentrated for immunodetection after RNAscope than for regular immunohistochemistry. The sections were mounted with Prolong Gold (Invitrogen, Cat: P36980) and DAPI for nuclei identification. These animals did not contain Fast Blue-labeled MNs; therefore, analyses were directed to MNs residing in known locations of the sciatic motor pool from L3 to L5 (N=4 sham (equal sex) and 5 sciatic nerve cut and ligation (2 male and 3 female). Animals were analyzed 14 days after sham or nerve surgery.

Microglia at all four locations (sham, contralateral, On MN, and DC) were imaged at 60x, 2x zoom, and 0.5 µm z-steps. Due to the thin tissue section thickness, entire microglia visualization within one section rarely occurs. Microglia were sampled for quantification if the cell body, nucleus, and at least a couple of processes were visible. *trem2* signal consisted of clumps of individual dots (each supposedly a single mRNA), which were frequently difficult to individualize. Therefore, in IMARIS, we obtained the total volume of *trem2* signal and normalized it to the “whole” eGFP microglia volume imaged or the DAPI nucleus volume. In death clusters, it was not possible to individualize microglia volumes, and therefore, *trem2* occupancy of cytoplasm and processes was calculated against the whole death cluster. Using this method n values for DC microglia were always smaller. DAPI-labeled nuclei in DC microglia were easily individualized and thus n is larger. Individual microglia outliers were removed via the ROUT method (Q=1%) in Prism (GraphPad). If any animal average was an outlier in a single analysis, the entire animal was removed from all analyses. Final sample sizes for statistical comparisons are in **Data S1** in support of respectively, **Figures 3B-C, 3D and 3E**.

**Figure 3.**
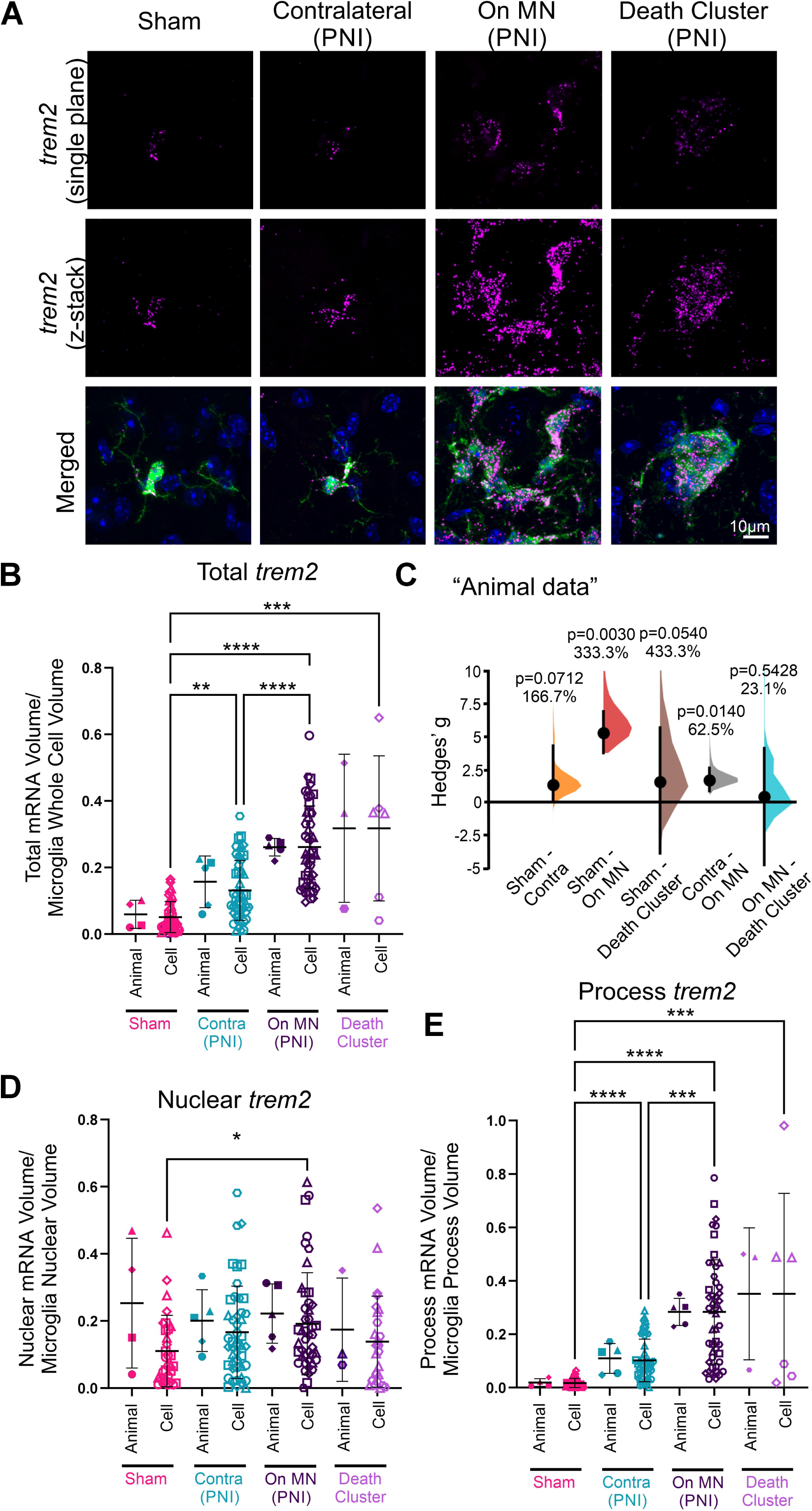
*trem2* mRNA expression is highest in activated microglia regardless of phenotype at 14 days post-injury. **A**) High magnification single plane confocal images showing *trem2* mRNA (magenta, RNAscope) in CX3CR1^eGFP^ microglia (green, eGFP) with DAPI (blue) delineating nuclei. Microglia, *trem2*, and nuclei were reconstructed in IMARIS and analyzed as volumetric 3D objects. **B**) Occupancy of total *trem2* mRNA signal in the eGFP microglia volume, including cell body, processes and nucleus. Individual microglia (Cell) and animal (Animal) averages are presented (n=10 microglia per animal per condition, N=4 sham (equal sex), 5 cut-ligation (2 males, 3 females). Each symbol represents one animal or microglia cell with symbol shapes identifying individual animals. Mean±SD is represented. For simplicity, only statistical comparisons from Dunn’s post hoc multiple comparisons for Cell data are shown (**p<0.01, ***p<0.001, ****p<0.0001) (see **Data S1** for further details). **C**) Total *trem2* comparison in animal data using estimation statistics. Black dots represent estimates of average Hedge’s g and vertical bars 95% confidence intervals. The estimated percentage change and significance from permutation t-tests are indicated in each comparison. Animals’ values for DC microglia bootstrapping were affected by a low n (n=3). All other comparisons confirm results in B. **D**) Analysis of *trem2* in nuclei delimited by DAPI. No significant differences were detected (Cell comparison: Kruskal-Wallis p=0.0501; Animal comparison: One-way ANOVA p=0.8716; **Data S1** for details). **E)** Same analyses as in B and D for *trem2* in microglia processes. The conclusions in processes mirror the total *trem2* data. Dunns comparison on Cells indicated **p<0.01, ***p<0.001, ****p<0.0001 (see **Data S1** for details).

To confirm *trem2* mRNA absence in GKO and CKO mice, we used the same methods, except that in this case, microglia were immunolabeled with IBA1 antibodies in GKO mice and genetically with tdTomato in CKO.

### P-Syk Immunofluorescence Analysis

We compared wildtype (Tmem119^eGFP^), GKO, and CKO animals that underwent sham or sciatic cut and ligation surgeries analyzed 14 days after surgeries. We considered the following experimental groups: Sham: N=4 WT (4 female), 6 GKO (4 male, 2 female), and 5 CKO (2 male, 3 female); PNI: N=6 WT (4 male, 3 female), 5 GKO (3 male, 2 female) and 7 CKO (4 male, 3 female). In this case, all animals contain Fast Blue-positive MNs retrogradely labeled from the LG muscle. Microglia were detected with eGFP (Tmem119^eGFP^ in WT and GKO) or tdTomato (Tmem119^CreERT2^ :: Ai9tdTomato in the CKO). eGFP was amplified with immunohistochemistry in its respective green channel. p-SYK was detected with phosphorylation-specific antibodies (see **Table 2**) in the Cy3 or Cy5 channel. Multiplex fluorescence sections were imaged with confocal microscopy (60x objective, 2x zoom, 1 μm step size), and 17-20 microglia per condition/genotype were randomly selected for quantification. The membrane of each microglia was manually traced in ImageJ, and the traced line width was set to 5 pixels (or ∼0.5 µm wide, images were approximately 0.1 µm/pixel). We measured the integrated pixel density under the line and under a similar line placed outside each cell to obtain a background estimate. Signal-to-noise ratios were calculated thereafter as explained for TREM2 immunofluorescence quantification. Individual microglia outliers were removed via the ROUT method (Q=1%) in Prism (GraphPad). If the animal average was considered an outlier, the entire animal was removed from the analysis. Final sample sizes are described in **Data S1** in support of **Figure 5B**.

**Figure 4.**
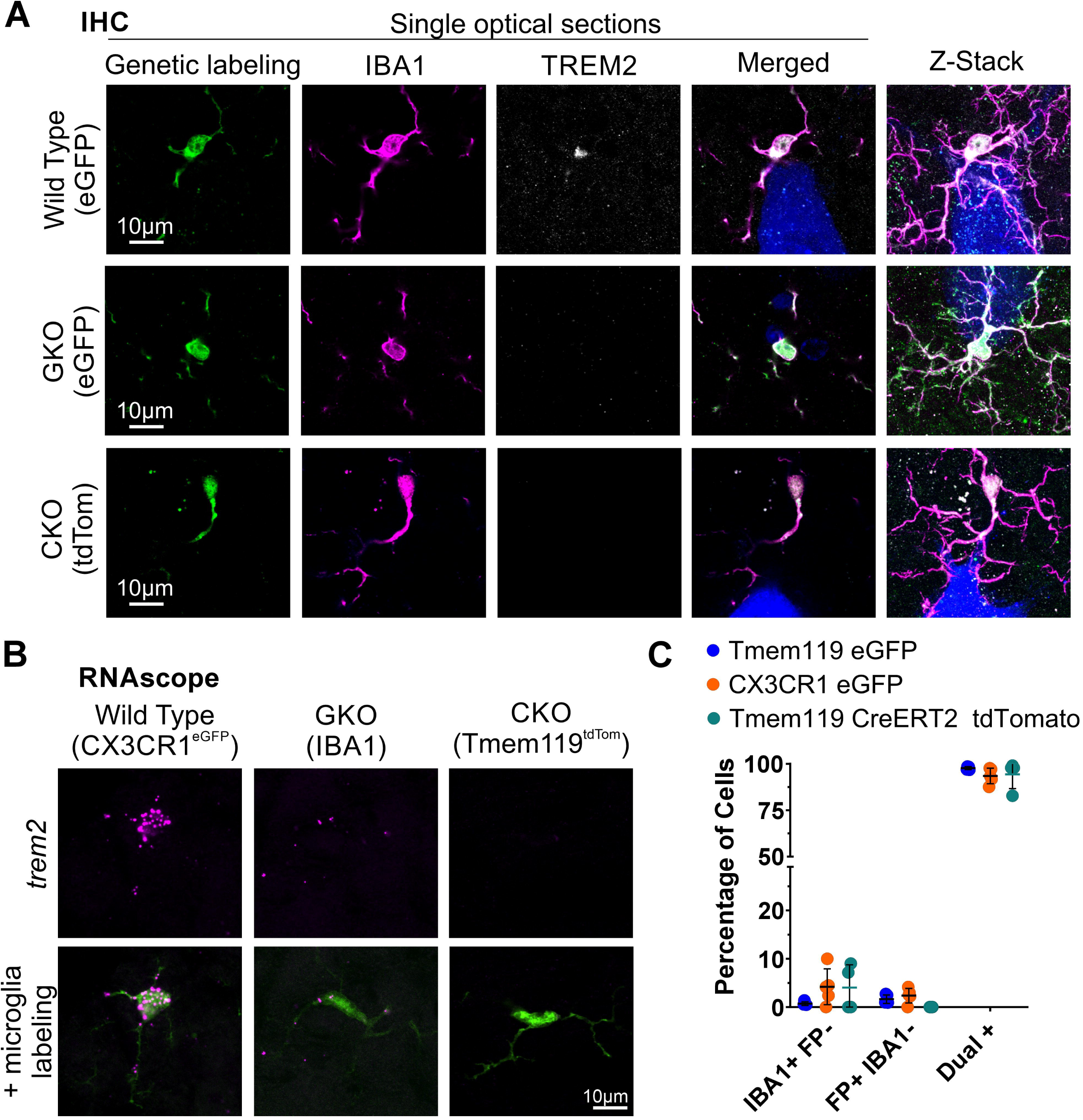
TREM2 global and conditional knock out validation. **A)** High magnification confocal images of TREM2 immunohistochemistry in microglia labeled either genetically (Tmem119 ^eGFP^ in WT and GKO or Tmem119^creERT2^:: Ai9 R26-lsl-tdTomato in CKO, all pseudocolored green for consistency) or with IBA1 (magenta). TREM2 (white) and Fast Blue (FB) motoneurons (MN) are also shown. TREM2 cytoplasmic and Golgi labeling are not present after either global (GKO) or conditional (CKO) deletion of *trem2*. **B)** High magnification confocal images of *trem2* RNA (magenta, RNAscope) and immunohistochemistry co-detection of microglia labeled by CX3CR1 eGFP in WTs, IBA1 in GKO, and tdTomato in CKO mice (all shown in green). mRNA *trem2* signal is absent in the CKO and greatly reduced in the GKO. **C)** Animal averages for the percentage of ventral horn cells with genetic fluorescent protein (FP) labeling and IBA1 in adult control Tmem119 ^eGFP^ (N=5 animals), CX3Cr1 ^eGFP^ (N=5 animals) and Tmem119^creERT2^:: Ai9 R26-lsl-tdTomato CKO animals (N=4). Theres is very good correspondence between genetic fluorescent protein labeling and IBA1 microglia; very Few cells are found with only genetic or IBA1 labeling. Recombination frequency in CreERT2 animals after tamoxifen is estimated 93.18% (n=5 L5 sections per animal, N=4).

**Figure 5.**
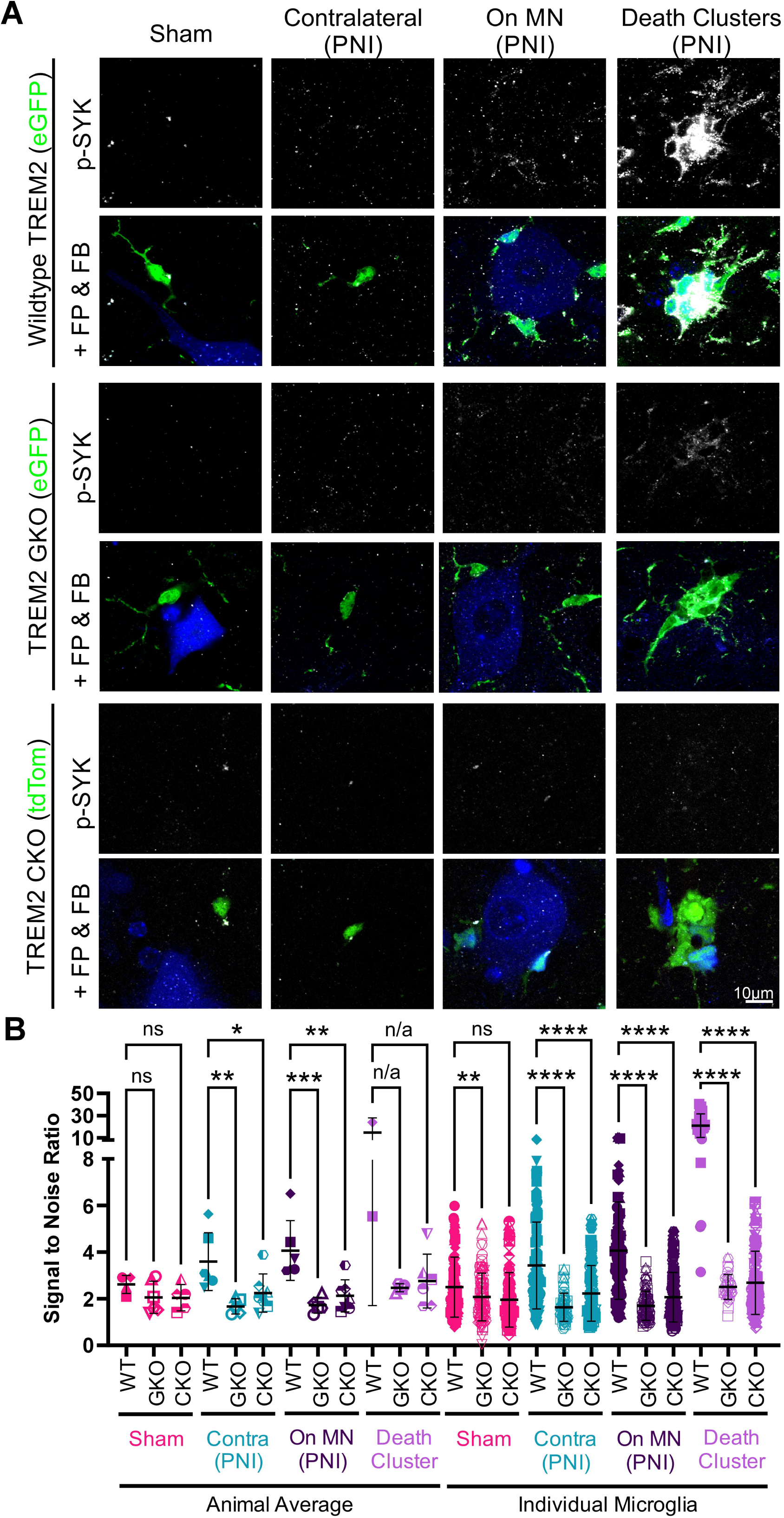
Phospho-SYK (p-SYK) increases with microglia activation 14 days post-cut-ligation and is diminished by TREM2 absence. **A**) High magnification single plane confocal images of microglia close to retrogradely labeled Fast Blue (FB) motoneurons (MNs) in sham mice or On MN and DC microglia in PNI mice. Microglia in the contralateral lamina IX of PNI mice are also shown. The top rows correspond to p-SYK immunolabeling (white), and the lower rows show in addition, genetically labeled microglia (green; FP: fluorescent protein) and FB MNs (blue). In wildtype (WT) and GKO microglia genetic labeling corresponds to Tmem119^eGFP^. In conditional KO (CKO) mice, genetic labeling is tdTomato (Tmem119^creERT2^:: Ai9 R26-lsl-tdTomato). For consistency, all microglia are pseudo-colored green. **B**) Quantification of p-SYK on microglia surfaces normalized to background (signal-to-noise ratio). Animal averages on the left (each data point is one animal average) and Cell averages on the right (each data point is one individual microglia). p-SYK is reduced in microglia from GKO and CKO mice, particularly in activated microglia. One-way ANOVAs followed by Bonferroni tests were used to compare animal averages. Individual microglia Cell data were compared by Kruskal-Wallis (p<0.0001) followed by Dunn test comparisons (n=17-20 microglia per condition per animal, N=4 WT female sham, 6 WT cut-ligation (equal sex), 6 GKO sham (4 males, 2 females), 5 GKO cut-ligation (3 males, 2 females), 5 CKO sham (2 males, 3 females), 7 CKO cut-ligation (4 males, 3 females). Significance for each pairwise comparison (Dunn tests): *p<0.05, **p<0.01, ***p<0.001, ****p<0.0001; no comparisons were performed for Death Cluster animal averages because of low n in WTs (n/a). Details can be found in **Data S1**.

### Ventral Horn Microglia Number and Territories in Sham Animals

We used N=7 WTs (3 male, 4 female), 7 GKO (4 male, 3 female), and 5 CKO mice (1 male, 4 female). Microglia were genetically labeled by eGFP (Tmem119^eGFP^ in WT and GKO) or tdTomato (Tmem119^CreERT2^:Ai9tdTomato in the CKO). eGFP was amplified with antibodies against its fluorescent protein. For each animal, we imaged five randomly chosen L5 sections with confocal microscopy and a 10x objective, 1x zoom, and 2 µm z-step. Only the ventral horn ipsilateral to the sham surgery was imaged. Analyses were performed in ImageJ on the collapsed z-stack confocal images (Jay et al., 2019). The ventral horn was outlined by tracing a horizontal line from the top of the central canal and outlining the gray-white matter border.

Within this ROI, individual microglia were identified and counted using the multi-point selection tool in Image J. The ROI area was measured, and a volume was obtained by multiplying by the section thickness (50 µm). From here, we calculated the average number of microglia within a volume cube of 100 µm side (10^6^ µm^3^) and obtained an average for each animal (n=5 sections per animal). Microglia territories were identified with the ImageJ Voronoi plugin and pseudo-colored for optimal definition. The average area of Voroni territories was calculated for each animal (n=5, L5 sections per animal). No outliers were identified via the ROUT method (Q=1%) in Prism (GraphPad). Exact sample sizes are reported in **Data S1** in support of respectively **Figures 6A and 6B**.

**Figure 6.**
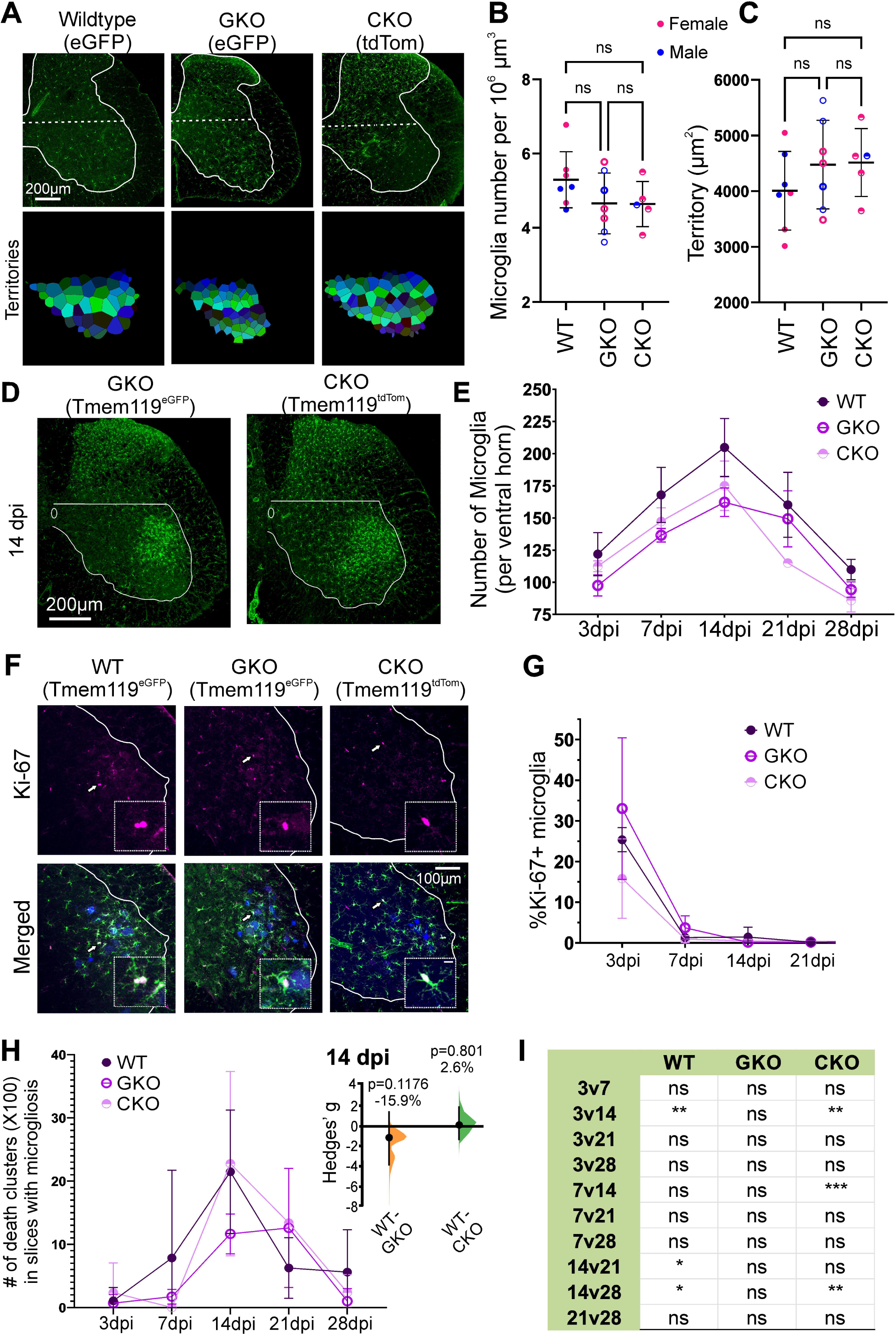
TREM2 knockout (KO) does not affect microglial numbers or their territories in controls or proliferation and death cluster formation after PNI. **A**) Low magnification collapsed z-stack confocal images showing spinal cord hemisections from Tmem119 ^eGFP^ “wildtype” (WT), Tmem119^eGFP^ TREM2 global KO (GKO), and conditional Tmem119^CreERT2^ td-Tomato microglia conditional KO (CKO) (microglia all in green for consistency). All animals had sham surgery. The grey matter is outlined in white and the border between the dorsal and ventral horn marked by a horizontal dashed lines (horizontal line from the top of the central canal). Each ventral horn microglia soma was counted, selected and used for Voronoi analysis in ImageJ to identify microglia territories shown below. The territory colors are arbitrary and only used to aid in individual territory distinction. **B**) Number of ventral horn microglia per ventral horn 100 µm side cube-volumes (10^6^ µm^3^). Similar numbers were found in WTs and TREM2 GKO or CKO. Each dot represents one animal. Mean±SD indicated. Each animal was estimated though five sections from the Lumbar 5 segment. N=7 mice for WT and GKO, N=5 for CKO. Pink, females; blue, males. One-way ANOVA failed to detect significance (p=0.2246). Post-hoc Bonferroni tests are indicated in the figure (ns=non-significant). **C**) Same analyses for territories. No significant differences were detected by one-way ANOVA (p=0.3068) followed by post-hoc Bonferroni tests (ns in graph). Details of statistical parameters in **Data S1**. **D**) Z-stack projections of confocal images from lumbar 5 sections (50 µm thick) 14 days after PNI of GKO and CKO mice with genetically labeled microglia (eGFP for WT and GKO and tdTomato for CKO, all shown in green). White outlines label the ventral horn and the central canal. **E**) Average number of microglia in the ventral horn at different times post-injury in WT, GKO, and CKO. Error bars=SEM. N=4 mice (equal sex) for each time point genotype, except 21 dpi CKO (N=2, both females). Each animal estimate was obtained from five Lumbar 5 sections. Two-way ANOVA revealed statistical significance according to time after injury (p<0.0001) and genotype (p=0.0157), although post-hoc Bonferroni tests did not detect significances across genotypes at any time point. Details in **Data S1**. **F**) Confocal images of immunolabeling for Ki-67 (white), microglia (green), and FB motoneurons (MNs) in WT, GKO, and CKO mice at 3dpi. White arrows point to Ki-67+ microglia examples featured in the zoomed-in images (white squares). **G**) Percent of Ki-67+ ventral horn microglia at different times post-injury in the three genotypes. Error bars=SD. Sample sizes as in A. Two-way ANOVA revealed significant differences according to time after injury (p<0.0001) but not genotype (p=0.1156). Details in **Data S1**. **H**) Number of death clusters at different post-injury times in WT, GKO, and CKO mice. The count represents all death clusters identified throughout the lumbar enlargement (Lumbar 1 to 6), divided by the number of sections analyzed and multiplied by 100. Error bars=SEM. Sample as in A. Two-way ANOVA detected significant differences according to time after injury (p<0.0001) with numbers peaking at 14 dpi, but not significant differences according to genotype (p=0.3989). Estimation statistics confirmed a lack of significant differences (Hedges’ g comparisons shown for the 14dpi time point). **I**) Results of Bonferroni post-hoc comparisons for time after injury in each genotype; *p<0.05, **p<0.001, ***p<0.001. Details in **Data S1**.

Time course of microglia proliferation and TREM2 expression in wildtype, GKO, and CKO These analyses were performed in animals that underwent a sciatic-nerve cut and repair injury to allow regeneration and analyze microgliosis at 3-, 7-, 14-, 21-, and 28-days post-injury (dpi). N=4 mice (equal sex) per condition (time after injury/genotype) except for CKO animals at 21 dpi, in which N=2 females. The proximal nerve stumps were soaked with Fast Blue in all animals to retrogradely label all axotomized MNs. Serial sections were immunolabeled with Ki-67 or TREM2 antibodies. Microglia were genetically labeled and fluorescent signals amplified (as before) with antibodies against eGFP (Tmem119^eGFP^ in wildtypes and GKOs). For microglia cell counts, we obtained confocal images from five random L5 sections with a 20x objective, 1x zoom, and 1 µm z-step. Microglia were manually identified and counted in Neurolucida. Animal averages were obtained from five L5 sections per animal. See **Data S1** in support of **Figure 6E**.

Ki-67 analysis was performed in the same sections. In this case, Ki-67 nuclear immunoreactivity was identified by a naïve observer blinded to the animal conditions, and a percentage of the total microglia detected was obtained for each section. Animals’ averages were obtained from five L5 sections per animal. Further analyses re-classified microglia according to location: On MN, in death clusters, in the white matter, and in the gray matter (different from On MN and DC microglia). Percentages of Ki-67+ cells for each microglia category were calculated. See **Data S1** in support of **Figure 6G**.

TREM2 quantification was performed in wildtype cut-repair animals following the same method explained above (“Analysis of TREM2 Immunofluorescence”).

### Number of Death Clusters comparison in wildtype, GKO, and CKO

We used the same time course mice to compare the number of death clusters at 3-, 7-, 14-, 21-, and 28-dpi. In addition, we compared the 14-dpi cut-repair time point with animals that underwent sciatic nerve cut and ligation surgery with no regeneration allowed. The data were compiled from animals used in previous analyses; N=12 WT, 12 GKO, and 8 CKO, all with equal sex numbers. An independent observer used a 20x objective on an Olympus BX60 epifluorescent microscope coupled to a RealTime Digital camera to count all death clusters throughout all recovered sections of the entire lumbar spinal cord in each animal. Only sections with microgliosis were counted. Quantification was blind to genotype and on serial sections to ensure death clusters were not double-counted or biased. The number of death clusters was divided by the number of sections to obtain a value per 50 µm-thick section in each animal. This value was multiplied by 100 to estimate the number of death clusters contained within 5 mm of rostro-caudal spinal cord length corresponding to the distribution of the lumbar region containing sciatic MNs from L4 to L6 segments. See **Data S1** in support of **Figure 6H**.

### Microglia morphologies in GKO mice

We reconstructed microglia morphologies for Sholl analyses as described before. N=6 animals and 5 microglia/animal reconstructed and skeletonized for each location: sham, contralateral, On MN and DC microglia (n=30 microglia per experimental condition). In this case, the nerve injury was a sciatic nerve cut and ligation, and the analyses were performed at 14 dpi. Microglia tracings underwent Sholl analyses (as above), and at each microglia location, data from GKO were compared to wildtype microglia data from **Figure 1**. See **Data S1** in support of **Figure 7B**.

**Figure 7.**
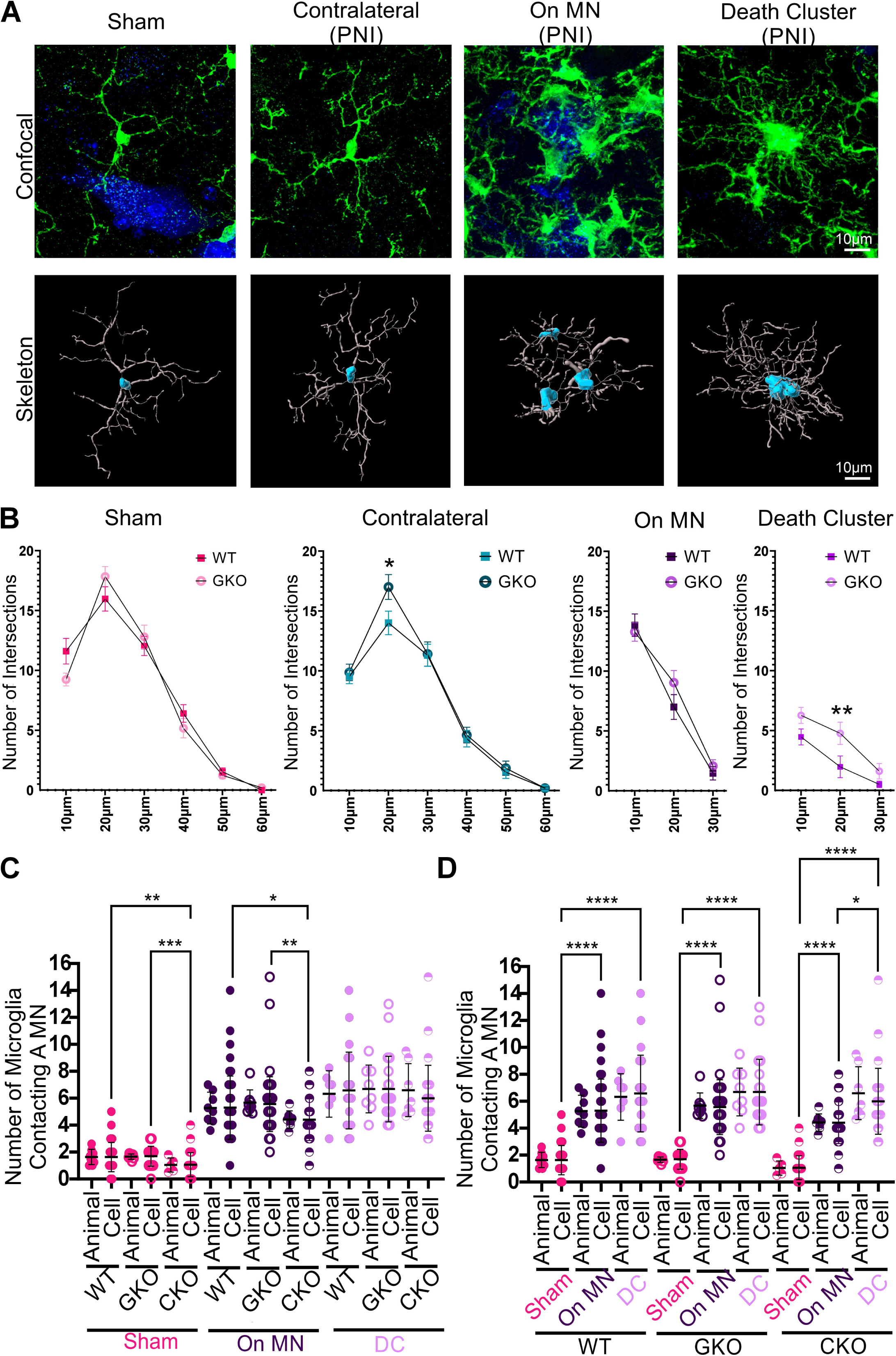
Morphology of activated microglia in TREM2 Global Knockout (GKO) 14 days post-cut-ligation. **A)** Top row: 2D projections of confocal image z-stacks of Fast Blue MNs and eGFP microglia (green) in either surveying (sham and contralateral) or activated states (On MN and DC microglia) in a GKO mouse. Bottom row: semi-automatically skeletonized microglia for Sholl analyses. **B**) Sholl analysis with WT data (from Figure 1E) showed small but significant differences in Contralateral and DC microglia in the 20 µm bin (*p<0.05, **p<0.01, Bonferroni post-hoc comparison WT vs GKO). Error Bars=SEM (pooled Cell data n=30 per genotype; n=5 microglia per condition per animal, N=6 mice (equal sex) per condition). Statistical details in **Data S1. C,D)** Number of microglia in death clusters or contacting MNs in sham or injured mice, organized by microglia type in **C** or genotype in **D**. Animal and Cell data mean±SD. N=8 sham and 8 injured in WTs, 7 sham and 8 injured in GKO, and 5 sham and 7 injured in CKO. 10 MNs analyzed per animal for sham or On MN microglia; after removing outliers n=75,77 MNs for sham and On MN in WTs, n=69, 67 in GKO, n=48, 68 in CKO. Death cluster sampling was n=33, 22, and 29 in WT, GKO, and CKO. One-way ANOVAs (Animal data) and Kruskal-Wallis (Cell data) showed significant differences according to microglia location in WT, GKO, and CKO mice (p<0.0001 in all cases, Dunn’s post-hoc comparisons for Cell data indicated in panel D). Comparisons across genotypes (panel C) revealed differences in sham and On MN in Cell data (p=0.0002, p=0.0017) but not in Animal data, which was underpowered (p=0.0635, p=0.0577). The number of microglia forming death clusters did not change in the three genotypes (p=0.9236 one-way ANOVA on Animal data and p=0.5729 Kruskal-Wallis on Cell data). Asterisk for Dunns pair-wise comparisons; *p<0.05, **p<0.001, ***p<0.001, ****p<0.0001. Further details in **Data S1.**

### Microglia-Motoneuron Interactions

The number of microglia contacting a MN or forming part of a death cluster were quantified by randomly selecting seven to ten Fast Blue-labeled MNs or up to ten death clusters per animal in 60x confocal images. To be selected, the entire MN and/or death cluster had to be visible within one confocal z-stack, which limited the total quantifiable MNs/death clusters. A microglia motoneuron contact was defined as an apposition of a microglia process or cell body on the surface of the MN cell body. The number of microglia with at least one contact was manually quantified by going plane-by-plane through the z-stack by a naïve observer blinded to experimental conditions and genotype. No outliers were identified via the ROUT method (Q=1%) using Prism (GraphPad). Final numbers of sampled MNs are reported in **Data S1** in support of **Figures 7C and 7D**.

### CD68 Quantification

We compared wildtype and GKO animals that underwent sham or sciatic cut and ligation surgeries. All animals were analyzed 14 days after the surgery. We analyzed the following experimental groups: Sham: N=8 WT (equal sex), 7 GKO (4 male/ 3 female); PNI: N=7 WT (4 male / 3 female), 5 GKO (4 male / 4 female). In all animals, MNs were retrogradely labeled from the LG muscle with Fast Blue. Microglia were detected with eGFP (Tmem119^eGFP^) amplified with immunohistochemistry. CD68 was detected with specific antibodies (see **Table 2**) in the Cy3 channel. The sections were imaged with confocal microscopy (60x, 2x zoom, and 0.5 μm step size) to sample sham, contralateral, On MN, and DC microglia and associated CD68 granules. We reconstructed in IMARIS 20-26 microglia per experimental condition (sham/injury, microglia type/location, and genotype) to calculate the eGFP-defined microglia volumes. We performed three types of estimates. The relative volume occupied by CD68 granules was calculated by extracting the total volumes of all objects with a CD68 signal detected in IMARIS and normalized to the total eGFP microglia volume and expressed as a percentage. In the case of death clusters, the total CD68 volume in the whole cluster was normalized to the whole death cluster volume since it was not possible to accurately separate individual microglia volumes in the death clusters. The number of CD68 granules was obtained from IMARIS segmentation of CD68 signals per microglial cell or death cluster. In the case of death clusters, the number of total granules was divided by the number of microglia composing the cluster to find a cell “average,” allowing a more appropriate comparison to individualized microglia. Maximum granule size was obtained via automatic detection in IMARIS in the image segmentation for each microglia or death clusters and measured in µm^3^. Individual microglia outliers were removed via the ROUT method (Q=1%) using Prism (GraphPad). If the animal average was considered an outlier, the entire animal was excluded. Final sample sizes are described in **Data S1 in support of Figure 8** (Sampling for CD68 Analyses), **Figure 8B** (CD68 analysis: Male Vs Female comparison in WTs), **Figure 9** data (CD68 total volume analysis, ANOVA analyses with GKO and CKO, total CD68 granule numbers analysis, maximum CD68 granule size analyses).

**Figure 8.**
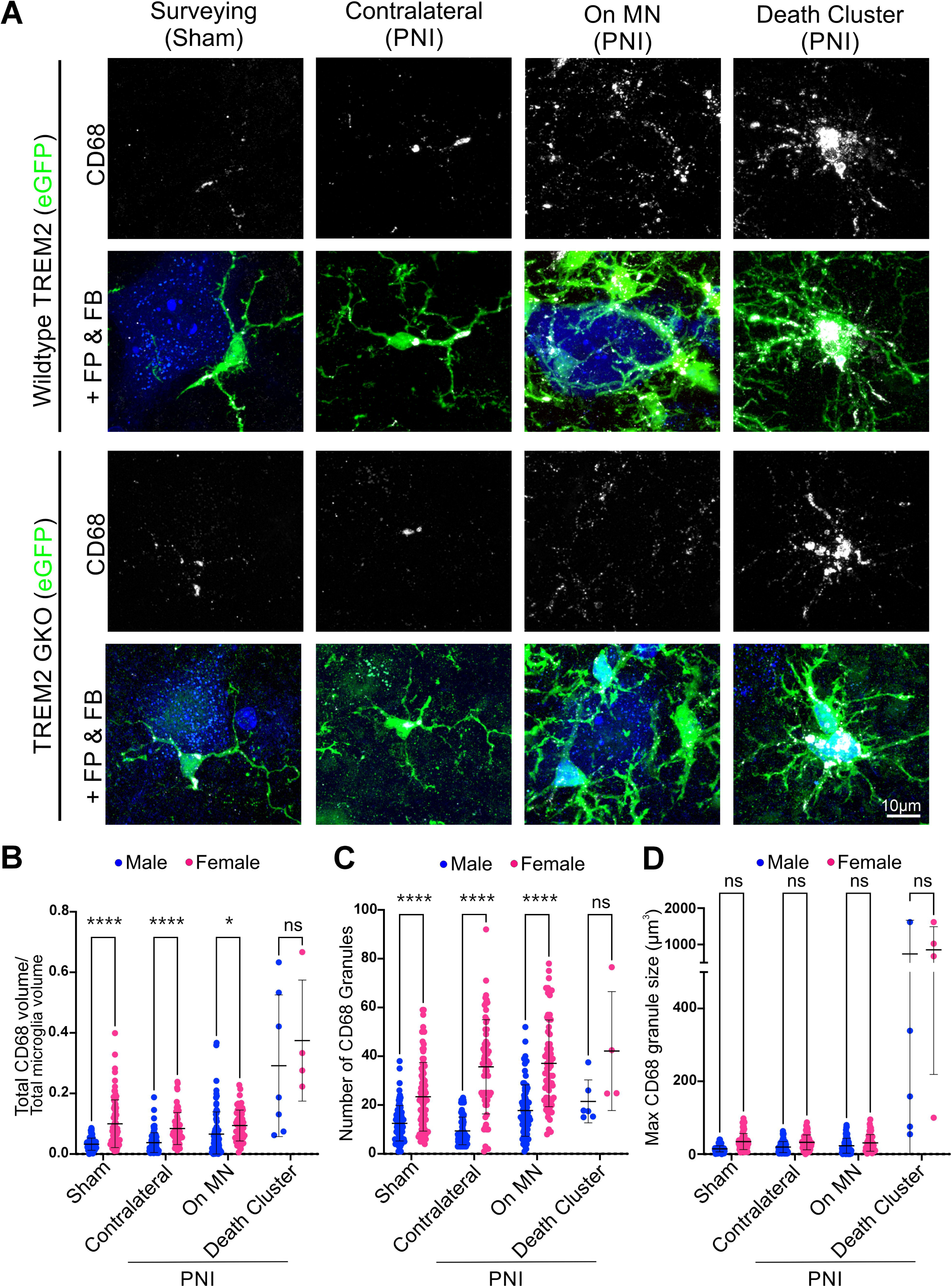
CD68 expression in control microglia and 14 days after injury; sex differences. **A)** High magnification confocal image z-stacks of microglia and Fast Blue (FB) MNs in WT and GKO mice after sham surgery or nerve injury (sciatic cut-ligation). Top rows: CD68 immunolabeling (white); lower rows added genetically labeled microglia (green FP: fluorescent protein) and FB+ MNs (blue). CD68 increases in activated On MN microglia, and it is maximal in DC microglia, being reduced in GKO mice. All images are from male microglia. **B,C,D**) CD68 expression sex differences. Sham surgeries mice: N=4 male and 4 female WTs and 4 males and 3 females GKOs. Nerve surgeries mice: N=4 male and 3 female WTs and 4 males and 4 females in GKO mice. 20-26 microglia sham, contralateral, and On MN analyzed per animal. After removing outliers n’s ranged from 61 to 91 for each experimental group. DC microglia recovered at lower rates with n’s between 4 and 13. Details in **Data S1**. **B**) Total CD68 to microglia volume ratios. Two-way ANOVA for microglia type and sex on all microglia pooled (Cell data) detected significance for both (p<0.0001). Bonferroni pair-wise tests significance indicated *p<0.05, ****p<0.0001 and confirmed by estimation statistics (**Data S1**). Except for DC microglia, females had higher CD68 content. **C**) Number of CD68 granules. Two-way ANOVA for microglia type and sex detected significance for both (p<0.0001). Bonferroni pair-wise test significance indicated, ****p<0.0001. Higher numbers of CD68 granules in sham, contralateral, and On MN female microglia were confirmed with estimation statistics. Sex differences in DC microglia did not reach significance with Bonferroni tests (p=0.0555) but did with estimation permutation t-tests (p=0.0452) (**Data S1**). **D**) Maximum size of the largest CD68 granule measured in each microglia. Two-way ANOVA for type of microglia and sex detected significance for microglia type (p<0.0001) but not sex (p=0.0504). Post-hoc Bonferroni comparisons did not detect significance (ns). However, estimation permutation t-tests detected small but significant differences with females expressing larger granules in surveying microglia (sham and contralateral, both p<0.0001, g=1.2 and 0.7 respectively) and On MN microglia (p=0.034, g=0.4), but not in DC cluster microglia (p=0.854).

**Figure 9.**
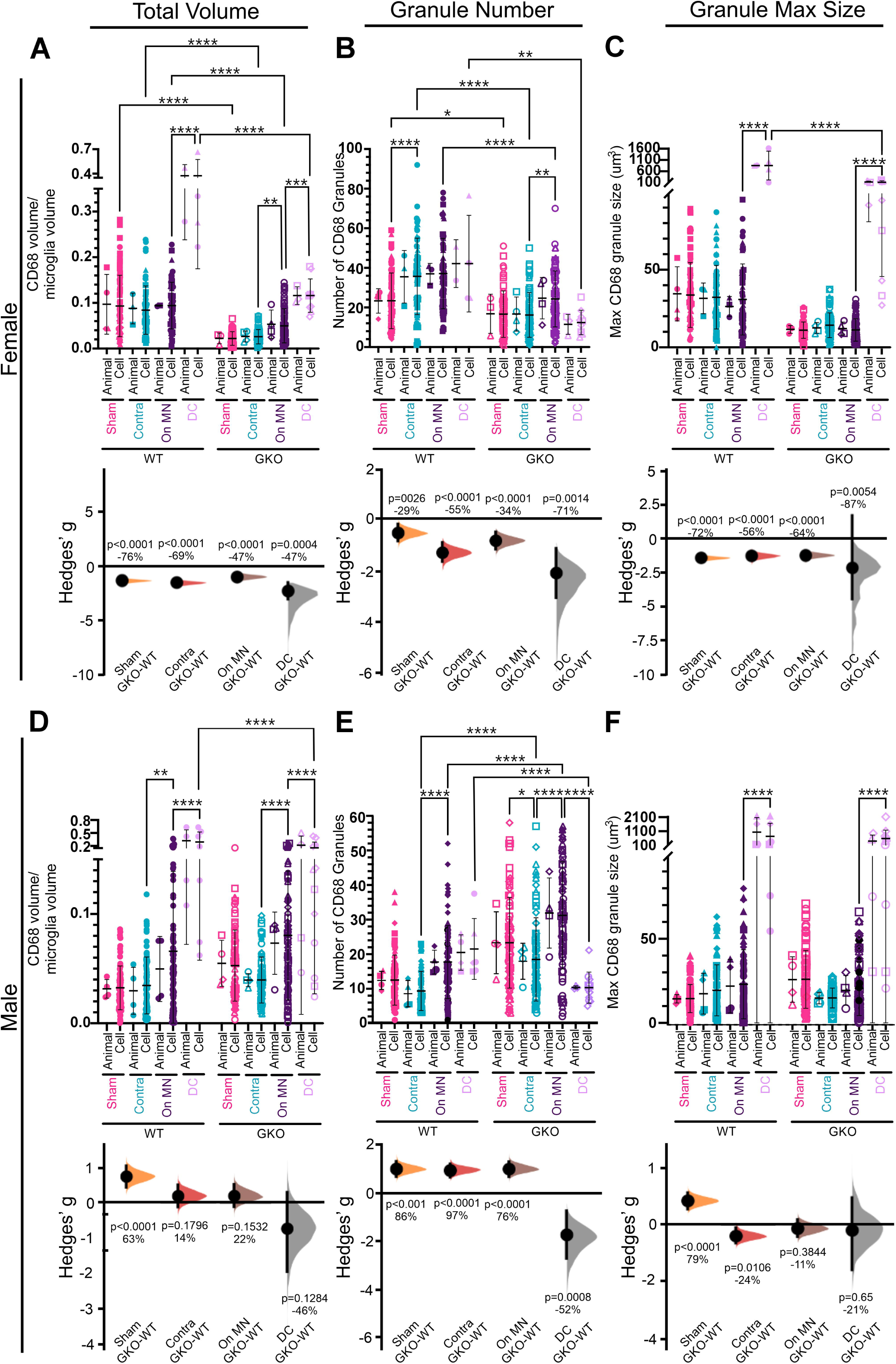
CD68 expression in different microglia types in wildtypes and TREM2 global knockouts in control microglia and after nerve injury. **A,B,C**) Female mice data for total CD68 volume (**A**), number of CD68 granules (**B**) and maximum CD68 granule size (*C*). The top graph shows the “Animal” and “Cell” data (mean±SD) and the results of two-way ANOVAs for type of microglia and genetics in the Cell data. Asterisks, *p<0.05, **p<0.001, ***p<0.001, ****p<0.0001 for post-hoc Bonferroni tests. Bottom graphs: estimation statistics, Hedge’s g estimates of mean differences between WT and TREM2 GKO in all four microglia categories represented by Cumming plots with bootstrap sampling distributions. Dots show mean g estimates; vertical error bars are 95% confidence intervals. Percentage differences and significance (permutation t-tests) are indicated on each graph. Sample summarized in **Data S1. D,E,F**) Same as above, but for male mice. Summary results: female TREM2 GKO mice show significantly lower CD68 expression compared to WTs in all three parameters studied (total volume, number, and maximum size of CD68 granules) and in all microglia: surveying (sham and contralateral) or activated (on MN and DC microglia). Male mice did not show consistent significant differences, except for the increase in the number of granules in TREM2 GKOs in sham, controls, and On MN microglia. Details of two-way ANOVAS, Bonferroni tests, Hedges’ g estimates, and permutation t-test are found in **Data S1** and the manuscript text.

### Analyses of motoneuron cross-sectional areas

To quantify the effects of TREM2 on cell body size changes during the chromatolytic reaction that occurs in response to axotomy we obtained 20x objective, 1x zoom and 1 µm z-steps confocal images of Fast Blue MNs at 3-, 7-, 14-, 21- and 28-dpi in wildtypes, GKO and CKO mice (N=4, 2 male / 2 female at all time points and genotypes, except for GKO at 28 dpi in which N=3, 1 male and 3 females and CKO at 21 dpi N=2 females, total N=57 mice). Confocal images were imported into Neurolucida, and an experimenter blinded to time after injury and genotype traced the outline of every cell body in the image in the optical plane in which the middle cross-section could be identified by finding the middle of the nucleus with an in-focus nucleolus. Five sections were measured per animal, and this represented a total of 7,090 manually traced MN cell body cross-sections or 124.4 ±28.9 (±SD) per animal with no significant differences among genotypes (WT: 118.3 ±26.0; GKO: 132.6 ±33.4; CKO 112.5 ±26.3). The area was measured in each traced Fast Blue positive MN cell body, and a distribution frequency histogram was calculated using 50 µm^2^ bin sizes for each animal. Animal averages for each genotype and time point were averaged to obtain one cell size distribution for each experimental condition (three genotypes X five time points = 15 conditions). Cell size distributions for each condition were then fitted to Gaussian distributions with n=2 (bimodal) using ClampFit (ver.11, Media Cybernetics). For statistical comparisons, we constructed cumulative-summation (cum-sum) distributions of the percentage of neurons placed in increasing size bins for each genotype across different times after injury. We also segmented the data for cells smaller than 450 µm^2^ (usual cut off for γ-MNs (Shneider et al., 2009) or larger than 1000 µm^2^ to detect any possible cell swelling at particular time points and genotypes. Further details of sample sizes and organization are in **Data S1** in support of **Figure 10D**.

**Figure 10.**
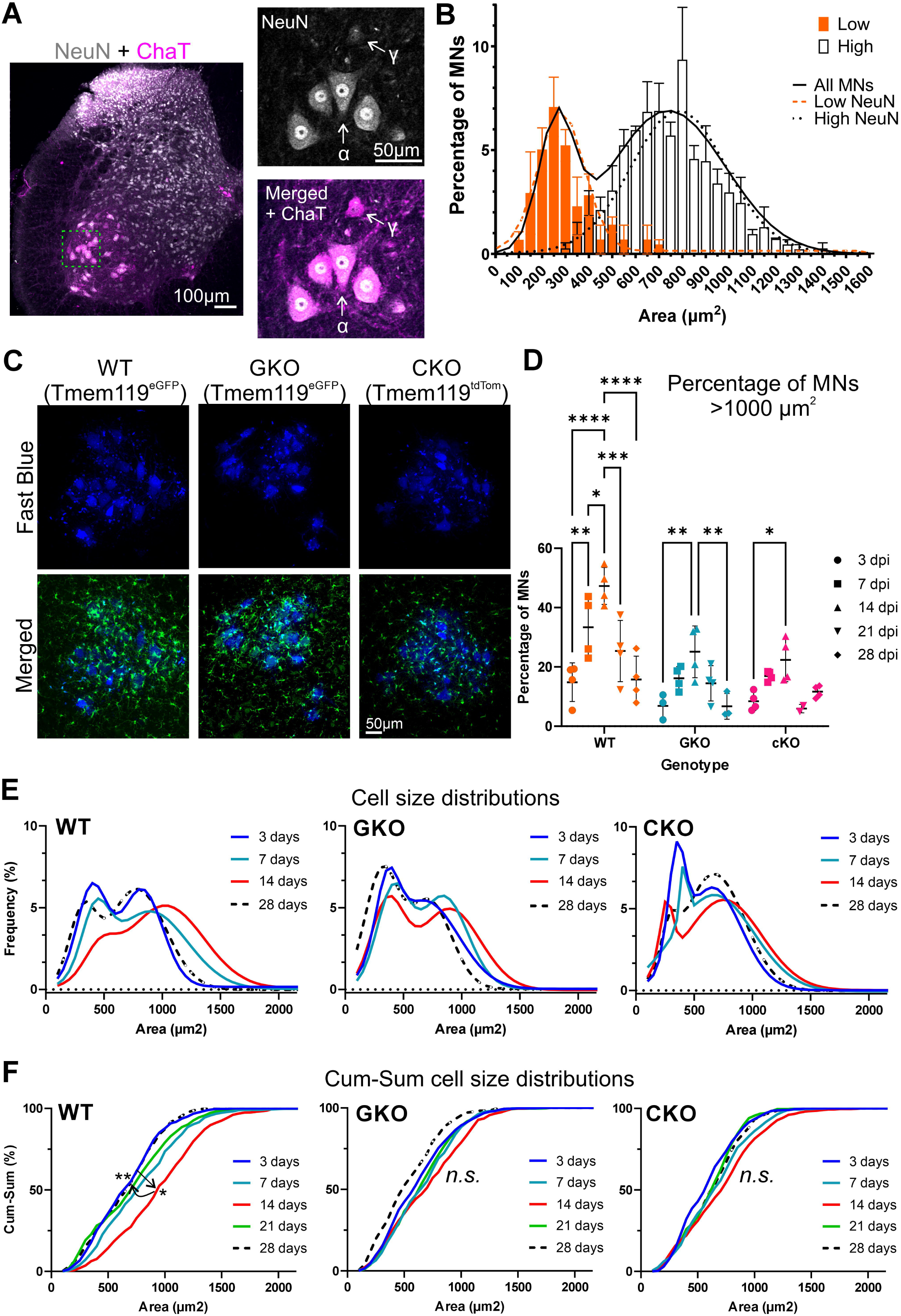
Effects of TREM2 deletion on the cell volume changes of regenerating motoneurons. **A**) Low magnification confocal image of NeuN immunoreactive neurons (grey) and ChAT-immunoreactivity (magenta). The boxed area on the left is magnified to show NeuN and NeuN merged with ChAT. α-MNs are ChAT+ and NeuN+; γ-MNs are ChAT+ and NeuN(-). **B)** cells distribution histogram in 50 µm^2^ bins (N=3 animals. Bars indicate mean ±SEM). Continuous lines are the bimodal Gaussian fit to all MNs (n=436 MNs). The dashed curves represent fits to low NeuN MNs (n=146) and high NeuN MNs (n=290). Data details in **Data S1**. **C)** Examples of the Fast Blue-labeled sciatic motor pools in WT, GKO, and CKO with microglia genetically labeled. **D**) Percentage of MNs with cross-sectional areas larger than 1000 µm^2^. The large increase noted at 24 dpi in WTs is blunted in GKO and CKO mice. Each dot represents one animal (in this analysis, we discarded one data point from one GKO at 3 days because it was an outlier). Mean percentage ±SD is represented. Two-way ANOVA for time after injury (dpi) and genotype showed significance for both (p<0.0001). Bonferroni pair-wise comparisons are shown for time after injury in each genotype. *p<0.05, **p<0.001, ***p<0.001, ****p<0.0001. Statistical details and comparisons between genotypes at each time point are in **Data S1. E**) Gaussian fits of cell size distributions of FB+ MNs at different time points in WTs, GKO and CKO. (n=437, 504, 475, and 419 MNs in WT at 3, 7, 14, and 28 dpi; n=435, 517, 563, and 429 in GKO, same time points; n=495, 467, 525, and 468 in CKO, same time points). There is a shift towards larger cells in WTs for 3 to 14 dpi, and a reduction in the γ-MN and α-MN peaks. Both return at 28 dpi. These changes are unclear in GKO and CKO mice. Details on fits are in **Data S1**. **F**) Cumulative plots of the same data compared with the Kolmogorov-Smirnov test. The shift towards larger values is apparent in WT data but not in GKO and CKO mice. Cell data distributions were significantly different only in the WT data between 3 days compared to 7 and 14 days (p=0.0150, 0.0282) and between 28 and 7 and 14 days (p=0037, p=0077). Details in Data S1.

To illustrate the significance of bimodal distributions in MN cell body size distribution, we analyze three naïve animals (1 female and 2 males) in which sections were immunostained for Choline Acetyltransferase (ChAT) and Neuronal Nuclear protein (NeuN). We measure all ChAT+ neurons in lamina IX of five L5 sections, focusing on the dorsal half of lamina IX corresponding to the sciatic motor pools (145.3 ±3.2 ±SD, MN cell bodies traced per animal). The traces were then divided into cells with high or low NeuN and broadly corresponding to α-and γ-MNs (n=96.7 ±11.8 high NeuN and 48.7 ±11.8 low NeuN ChAT+ cell bodies traced per animal). The data were fitted to Gaussian distributions as above for the whole MN population and each of the high and low NeuN ChAT+ MN populations. Further details of sample sizes and organization are in **Data S1** in support of **Figure 10B**.

To analyze MN cell size distributions after regeneration and muscle reinnervation 8 weeks after the nerve injury we further analyzed wild types either Sham controls (N=6, 3 males and 3 females; n = 1,286 MNs or 214.3 ± 54.5 per animal ±SD) or with a sciatic nerve cut and repair injury (N=3 females; n = 464 MNs or 154.7 ± 63.7 per animal ±SD). We compared the cell size distribution in Sham and injured wildtypes ipsilateral to the injury to both the ipsilateral (injured-experimental, n=243 and 254 MNs respectively) and contralateral (uninjured-control, n=209 and 250, respectively) sides of CKO animals (N=2, 1 male and 1 female). We immunostained for ChAT and NeuN with similar procedures as above and followed the same analytic procedures and curve fitting protocols.

### Analysis of neuromuscular junction (NMJ) innervation

Confocal images (20x objective, 1x zoom, z-step=1.5 µm) were taken with an Olympus FluoView FV1000 confocal microscope. Thereafter, LG muscle neuromuscular junctions (NMJs) were analyzed manually in ImageJ, blind to surgery condition and sex. Max projection z-stacks were collapsed and transformed into a binary image via manual thresholding of each channel. A ROI was drawn around 50 *en face* NMJs per animal. The area of Vesicular Acetylcholine Transporter (VAChT) and α-Bungarotoxin (α-BTx) was measured to acquire the percentage of innervation area *per* NMJ. The VAChT signal was always less than the α-Bungarotoxin signal such that in controls >50% coverage defined fully innervated NMJs. The percentage of fully innervated NMJs was calculated per animal. Sample sizes and organization are in **Data S1** in support of **Figure 12**.

**Figure 11.**
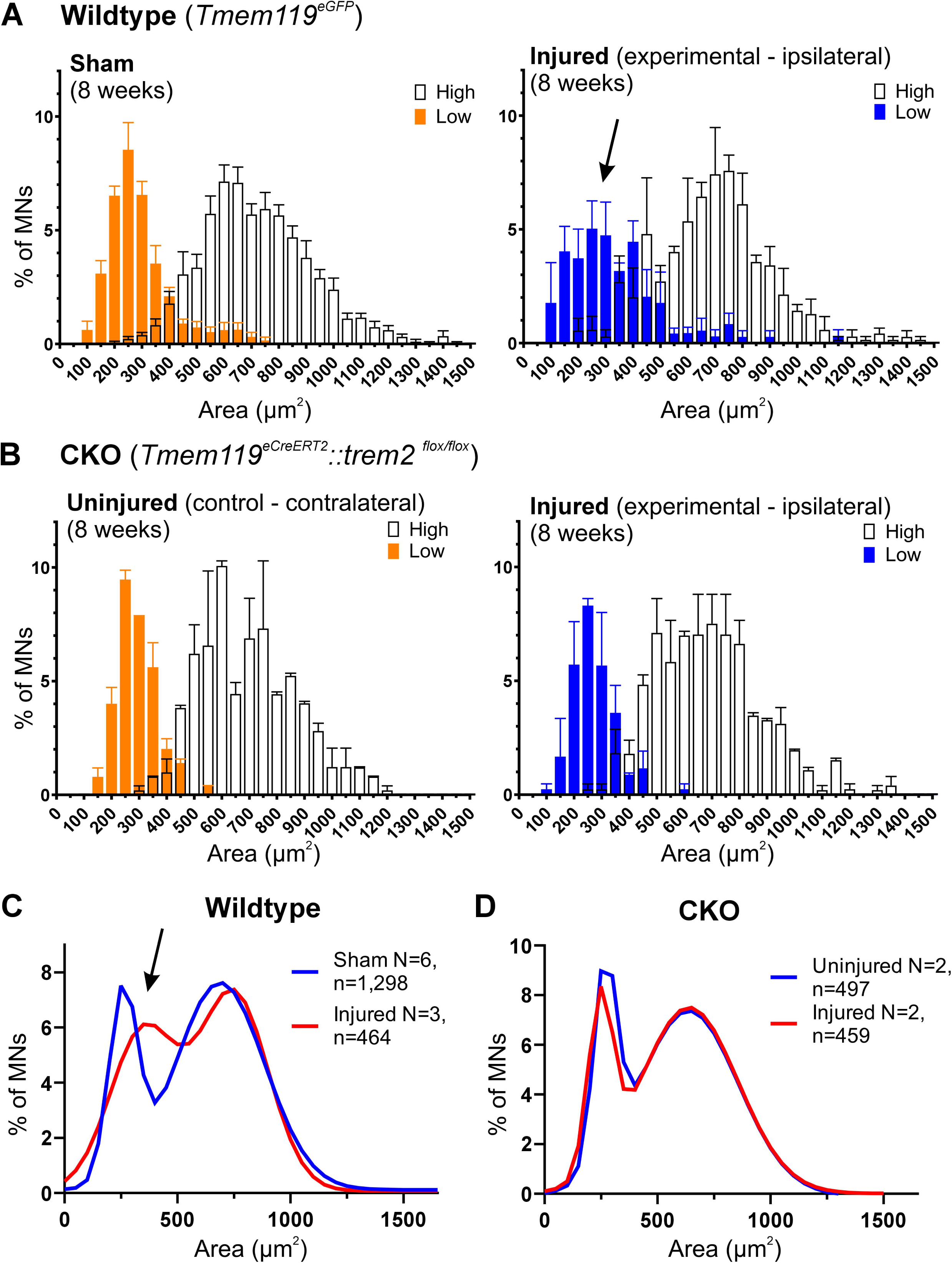
Deletion of TREM2 in microglia rescues a subpopulation of small size MNs, possibly. γ**-MNs**. **A**) Cell size distributions of MNs and correspondence with high and low NeuN content in wildtype animals 8 weeks after a sham surgery (left graph) or a sciatic nerve cut and repair injury (right graph). The data is distributed in 50 µm^2^ bins. Sham animals N=6 (equal sex), Injured animals, N=3 (all females). Each bar shows the average percentage of MNs with cell sizes within each bin and error bars indicate SEM. The small size peak with low NeuN is usually interpreted as composed mainly of γ-MNs. Eight weeks after injury a proportion of MNs are missing in this peak (arrow). Additionally, many NeuN-negative MNs are in size bins intermediate between the two peaks. The large sized peak of MNs with high NeuN content is interpreted as α-MNs. **B**) MN cell size distribution histograms and NeuN content as in A, but in CKO animals 8 weeks after a cut-repair injury of the sciatic nerve (N=2, 1 male and 1 female). Graphs show the uninjured control side contralateral to the injury (left graph) compared to the ipsilateral side containing injured MNs after 8 weeks of regeneration (right graph). The small size peak of MNs with low NeuN content and corresponding to γ-MNs is preserved in CKO animals. **C**) Cell size distribution of all MNs sampled in each experimental group and fitted with binominal distributions in sham control MNs (blue; n=1,286 MNs) and injured/regenerated MNs (red; n=464). The arrow points to the small size peak showing decreased amplitude and a shift towards large size bins 8-weeks after injury and in the regenerative period. **D**) Same cell size distributions and binomial fitting but for MNs in CKO animals sampled from the control side contralateral to the injury (blue; 497 MNs) and the experimental side ipsilateral to the injury (red; 459 MNs). Compared to panel B there is preservation of the γ-MN peak. Both MN cell size distributions appear identical to each other in the absence of microglia TREM2 activity.

**Figure 12.**
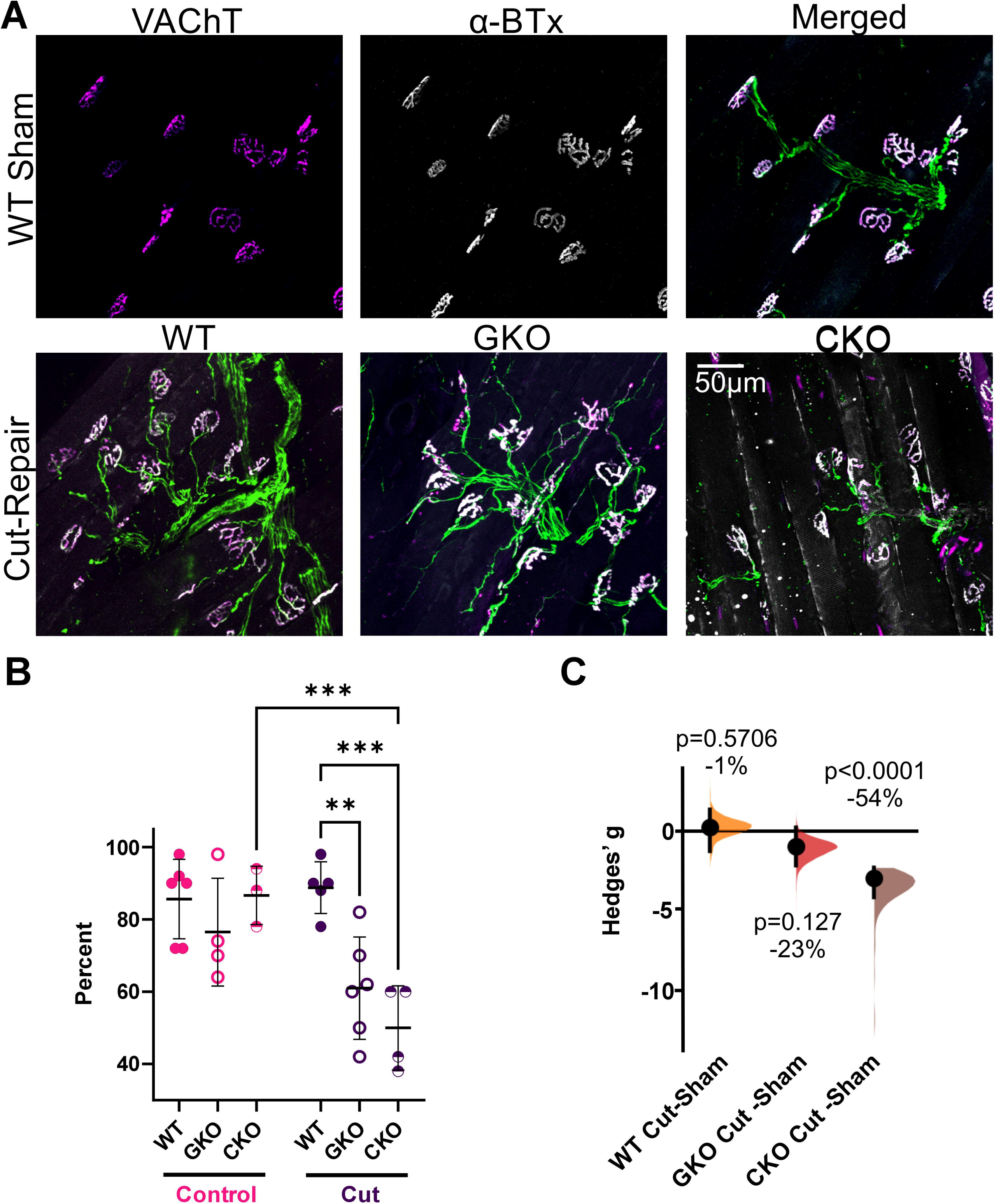
TREM2 deficiency reduced neuromuscular junction (NMJ) reinnervation eight weeks post sciatic cut-repair. **A)** Low magnification (20x/0.70 NA) confocal images of lateral gastrocnemius (LG) muscles with immunohistochemistry for Vesicular Acetylcholine Transporter (VAChT, red) (labels axon terminals), α-Bungarotoxin (α-BTX, blue) (labels motor end plate), and Neurofilament Heavy Chain (NFH, green). **B)** The percent of fully innervated NMJs analyzed via Two-Way ANOVA with Bonferroni’s post hoc multiple comparisons (each data point is one animal, n=50 per mouse, mean ±SD are represented, WT vs GKO p=0.002, WT vs CKO p=0.0002, CKO control vs cut p=0.0005). **C)** NMJ innervation analyzed with estimation statistics. Hedges’ g (Ho et al., 2019) of mean differences represented by Cumming plots with bootstrap sampling distributions. Dots show means; vertical error bars are 95% confidence intervals. Percentage differences and significance (permutation t-tests) are indicated on each graph. Statistical details in **Data S1**.

### Statistics

Each dataset was analyzed for normality using Shapiro-Wilk tests. Normally distributed samples were compared using parametric statistics: one-way or two-way ANOVAs, followed by Bonferroni-corrected pair-wise multiple comparisons. Non-normally distributed samples were compared using Kruskal-Wallis ANOVA followed by Dunns’ test pair-wise multiple comparisons. In a few cases, we compared only two data sets. In this case we used two-tailed t-tests if data were distributed normally or non-parametric Wilkinson tests if normality failed. For each dataset, we performed statistical comparisons for “Animal” and “Cell” data. “Animal” data consisted of comparing the averages of different animals, in which each animal is considered an independent data point. “Cell” data consisted of pooling all the data (microglia cells or MNs) from all animals in the same experimental group. Each type of comparison has advantages and disadvantages when describing the results (Dukkipati et al., 2017). “Cell” data preserves the original biological variability and has a large n, reducing sample size errors. “Animal” data loses the original biological variability and results in smaller n, thus inducing Type I errors, particularly when n is small, for example, when sex differences are taken into consideration, or when many multiple comparisons are performed simultaneously. But in “Animal” data all data points are independent while in “Cell” data, not all values are independent (repetitive measures in the same animal). We minimize this problem by obtaining similar sample sizes and repetitions in all animals for each statistical comparison. In our data sets, “Cell” data always had higher variability and was always skewed and non-normally distributed, while “Animal” data was always normally distributed with lower variability. Statistical comparisons of both datasets were performed in parallel, and significance was set at p<0.05. In general, there was good agreement between both, and when this was not the case, it is noted in the results. Throughout the text, N refers to the number of animals analyzed and n to the number of cells/sections.

In addition, we confirmed statistical conclusions by estimating effect sizes using estimation statistics (Ho et al., 2019). In this case, we calculated mean differences and Hedge’s g for each pair-wise comparison and the 95% confidence intervals of estimates performing 5000 bootstrap subsamples with recalculation of both parameters. This allows the creation of Gaussian distributed mean differences and Hedges’ g estimates, making comparisons independent of original distribution properties on normality and variance. The data are represented by Cumming plots with bootstrap sampling distributions of estimates and dots representing the average estimate and vertical bars the 95% confidence intervals. The null hypothesis is that the difference between the samples equals 0. Rejection of the hypothesis occurs when 95% confidence limits do not cross 0 (less than 0.05 probability), in which case we conclude the calculated effect size is significant and the samples are different. This is further confirmed with permutation t-tests with p<0.05. In general, there was good agreement between parametric, non-parametric, and estimation statistics in the different “Animal” and “Cell” samples, although estimation statistics tended to be more sensitive for detecting significance when n was small or when the number of multiple comparisons performed generated corrections in post-hoc comparisons that significantly increased Type I errors. We investigated possible sex effects with two-way ANOVAs for sex and conditions (i.e., sham/PNI; genotype, microglia types) to detect possible interactions. Sex comparisons are reported in supplementary figures, except when sex differences are critical for interpretation, in which case they are included in the main manuscript figures. To compare cell size distributions, we used cum-sum plots and Kolmogorov-Smirnov (K-S) statistics. We compared differences in cell size distributions at different survival times within each genotype (WT, GKO and CKO).

Parametric and non-parametric statistics were performed in Prism (GraphPad), cell distribution fits and K-S tests were performed in Clampfit (Media Cybernetics), and estimation statistics in the web app (https://www.estimationstats.com/<u>#/</u>). All statistical parameters, sample sizes, and effect sizes calculated are fully reported in a supplementary **Data S1** excel file contains all statistical Tables (one tab for each analysis).

## Results

### Activated microglia display two morphologically distinct interactions with injured motoneurons 14 days post sciatic nerve transection

To study microglia-MN interactions, we used a Tmem119^eGFP^ reporter mouse in which microglia express enhanced Green Fluorescent Protein (eGFP) without interfering with *Tmem119* endogenous gene expression (*Tmem119-2A-GFP* (Kaiser & Feng, 2019)). For identification of injured MNs, a subset of sciatic nerve MNs were retrogradely labeled with Fast Blue (FB) from the lateral gastrocnemius (LG) muscle five days before nerve surgeries. Animals were analyzed 14 days post-injury (dpi), corresponding to the peak of the microglia reaction in the ventral horn after PNI (Rotterman et al., 2019; Rotterman et al., 2024) (**Figure 1A**). Microglia eGFP fluorescence was amplified with antibody staining to optimally reveal morphological details. Spinal cords from sham animals contained typical surveying microglia with processes that occupied largely non-overlapping territories. After PNI (in this case, a sciatic nerve cut and ligation), there was a prominent microglia reaction in the ventral horn region around the injured sciatic MNs with microglia migrating towards the MN cell bodies (**Figure 1B**). Some microglia attached to the cell bodies of MNs, with two distinct microglia-MN interactions (**Figure 1C**) (Rotterman & Alvarez, 2020). More frequently, microglia remained individualized from one another while they partially enwrapped the cell body of MNs with smooth MN cell body edges and no evidence of degeneration. We refer to this phenotype as “On MN” microglia. A second interaction was less frequent and consisted of ameboid microglia that tightly cluster together, forming what we named a “Death Cluster” because it fully surrounds degenerating MNs with irregular and warped cell bodies.

To generate a quantitative description of these different morphologies, microglia were semi-automatically reconstructed using IMARIS software (**Figure 1D**) and Sholl analyses performed after skeletonization (**Figure 1E**). On MN and Death Cluster (DC) microglia were compared to surveying ventral horn microglia sampled from sham animals or the contralateral ventral horn (n=5 microglia per animal; N=6 animals per condition, equal sex numbers). No sex differences were detected (**Figure S1**), and male and female data were pooled together. Surveying microglia had ramified processes extending as far as 60 µm from the microglia cell body. On MN microglia were also ramified, but their processes were shorter, hardly extending beyond 30 µm from the cell body. DC microglia were the least ramified, with fewest and shortest processes (**Figure 1D-E**). Two-way ANOVA for Sholl distance and microglia location revealed significant differences among the four experimental groups (p<0.0001 for both variables and their interaction, statistical details in **Data S1** in support of **Figure 1E**). Post-hoc pair-wise comparisons (Bonferroni tests, **Figure 1E**) demonstrated that Sholl intersections were not significantly different between sham and contralateral control microglia, but both were significantly different from On MN and DC microglia (**Figure 1** and **Data S1** in support of **Figure 1E**). Compared to sham, the number of intersections in On MN and DC microglia was reduced by 56% and 88%, respectively at 20 µm distance and by 89% and 96% at 30 µm distance, with a negligible number of processes reaching the 40 µm distance bin. DC microglia showed fewer processes close to the cell body (10 µm Sholl bin) when compared to any other microglia group (p<0.0001 in all cases). On MN microglia have significantly longer and more complex processes 20 µm from their cell body compared to DC microglia (p<0.0001). In summary, microglia morphologies are distinct depending on the health status of the MN they associate with, being more phagocytic-like in DC microglia.

### TREM2 expression increases in all activated microglia after PNI but Death Cluster microglia contain the highest expression

Cellular mechanisms relevant for regeneration or cell death of axotomized MNs like neuronal phagocytosis, synaptic pruning and metabolic homeostasis have been associated with microglial TREM2 in other neuropathological contexts (Filipello et al., 2018; Hsieh et al., 2009; Jay et al., 2019; Qu & Li, 2020; Rueda-Carrasco et al., 2023; Scott-Hewitt et al., 2020; Tagliatti et al., 2024; Takahashi et al., 2005; Yu et al., 2023). Therefore, we analyzed TREM2 protein expression on Tmem119^eGFP^ labeled activated microglia associated with injured MNs of different fates (**Figure 2A**). TREM2 immunofluorescence intensity on the cell membrane and cytoplasm (excluding the intense patch in the Golgi (Sessa et al., 2004)) increased in On MN microglia and was highest in DC microglia (**Figure 2A**). Increased immunoreactivity outlining microglial cell bodies suggests TREM2 translocation to the plasma membrane. Quantitative comparisons were performed with all microglia pooled together (“Cell”; n=160 sham; n=154 contralateral; n=181 On MN; n=52 DC) or on animal averages (“Animal”; N=8 sham and 8 PNI animals, equal sex). Male and female data were pooled together because no sex differences were found in TREM2 expression (**Figure S2**). “Cell” data preserved the original sample cell variance, and distributions were always skewed towards high values in a proportion of microglia. “Animal” averages dampened cell-to-cell variations and were always normally distributed. Significant increases in expression were detected in activated microglia in both data sets **(Figure 2B**, **Data S1** in support of **Figures 2B-C**). “Cell” data for On MN and DC microglia showed respectively 78% (effect size Hedges’ g=1.7) and 125% (g=2.1) more TREM2 compared to sham microglia (p<0.0001, Kruskal-Wallis ANOVA followed by Dunn’s post-hoc multiple comparisons). DC microglia showed 62% more TREM2 than On MN microglia (p=0.002, Dunn’s test; g=1.2). Estimation statistics of bootstrapped mean differences followed by permutation t-tests confirmed these conclusions (**Data S1** in support of **Figures 2B-C**). “Animal” data replicated the significant increases in DC microglia compared to sham or On MN (p<0.0001 and p=0.0125, respectively; Bonferroni tests), and this was confirmed with estimation statistics that also uncovered the difference detected in “Cell” data between sham and On MN microglia (p=0.0012, permutation t-tests, **Figure 2C**). Despite a small TREM2 increase in contralateral microglia the differences with sham microglia did not reach statistical significance (**Figure 2C**, **Data S1**).

Next, we analyzed the time course of TREM2 expression after nerve injury. To best interpret the data in the context of regeneration we performed nerve injuries followed by repair with fibrin glue (Akhter et al., 2019) and analyzed animals at 3, 7, 14, 21, and 28 days post-injury (dpi) (**Figure 2D**) (N=4 animals at each dpi, n=20 microglia analyzed per animal for On MN and contralateral; after removing outliers n varied between 72 and 79 microglia in different groups, see **Data S1** in support of **Figure 2E**). DC microglia numbers were variable at different times post-injury; they were rare at 3 dpi, scarce at 7 and 28 dpi, and frequent at 14 and 21 dpi. This suggests that most MN cell death/removal occurs during the second and third week after injury. The different rates of death cluster formation at different time points created variability in the number of microglia (n) and animals (N) analyzed for DC microglia (3 dpi n=16, N=1; 7 dpi n=29, N=2; 14 dpi n=80, N=4; 21 dpi n=48 N=3; 28 dpi n=37, N=2). Since the number of animals with recoverable data in death clusters was low at some time points, the data from different animals were pooled into “Cell” data sets for analyses. TREM2 expression according to time after injury and microglia type (contralateral, On MN and DC) showed significant variations according to both variables (two-way ANOVA, p<0.0001, **Figure 2E**, **Data S1**). To reduce Type II statistical errors, we limited pair-wise comparisons (Bonferroni) to adjacent dpi data sets within the time course of each microglia type (contralateral, On MN, and DC). In follow-up analyses we compared all three microglia types at each time point. Salient findings include a significant increase in TREM2 expression at 14 dpi for On MN and DC microglia compared to 7 dpi (p<0.0001 in both cases, 68% and 64% increases) that thereafter significantly decreases in microglia On MN at 21 dpi (14 to 21 dpi, p=0.0083; 25% reduction) and is further reduced at 28 dpi (21 to 28 dpi, p=0.0015; 37% decrease) (**Figure 2E**). In contrast, DC microglia maintain TREM2 expression at 21 dpi (14 to 21 dpi, p>0.9999) and this is partially decreased by 28 dpi (21 to 28 dpi, p=0.0035; 24% decrease) (**Figure 2E**). Comparisons among different microglia types at each time point revealed that TREM2 expression in DC microglia is higher at all time points compared to either contralateral (between 66% and 319% increase at different time points) or On MN microglia (63% to 141% higher), while On MN microglia TREM2 expression was significantly higher than contralateral microglia at 14 dpi (96% higher) (Details in **Data S1** in support of **Figure 2E**). In conclusion, TREM2 expression parallels the time course of the microglia reaction, peaking at 14 dpi ipsilateral to the injury. DC microglia upregulate TREM2 at higher levels throughout the period of microglia activation (from 3 to 28 dpi) and retain expression for longer after the peak.

To investigate whether changes in TREM2 protein were associated with changes in *trem2* gene expression, we evaluated *trem2* mRNA using RNAscope in CX3CR1^eGFP^ mice at 14 dpi after sciatic nerve cut and ligation (**Figure 3A**). We used CX3CR1^eGFP^ mice because endogenous fluorescence withstood quenching from the RNAscope protocol unlike in Tmem119^eGFP^ mice. Over 90% of IBA1 detected microglia were labeled in both CX3CR1^eGFP^ and Tmem119^eGFP^ mice with similar specificity, suggesting both models label the same microglia with similar sensitivity (**Figure 4C**). Robust *trem2* upregulation in activated microglia resulted in clumping of the granular signal, preventing quantification of individual granules. We therefore estimated the occupancy of *trem2* fluorescence within microglia volumes delineated by eGFP or the nucleus marked with DAPI (see methods) (n=35 sham; n=47 contralateral; n=49 On MN; n=6 DCs; N=4 sham equal sex, and N=5 in PNI animals, 2 male/3 female). It was not possible to individualize microglia in death clusters for IMARIS 3D reconstructions. In this case, we obtained an average occupancy ratio of *trem2* mRNA per whole microglia cluster. This explains the lower n in DC microglia data. A sex difference was found in total *trem2,* being higher in female DC microglia vs male (p=0.026, Bonferroni, **Figure S3**). We pooled data from both sexes to overcome the low number of sex-specific death clusters and because no differences were found in other microglia. Like the TREM2 protein, *trem2* mRNA was evaluated as “Cell” or “Animal” data (**Figure 3B**). Compared to sham, On MN and DC microglia had significantly higher total *trem2* mRNA, whether compared in “Cell” or “Animal” data (Cell: p<0.0001, Kruskal Wallis; Animal: p=0.0173, one-way ANOVA). Also, like protein analyses, effect sizes were large. On MN microglia showed 420% and 333% increases in *trem2* mRNA when compared to sham by “Cell” (g=2.1, p<0.0001, Dunns or permutation t-test) or “Animal” data (g=5.3, p=0.0519, Bonferroni; p=0.0330, permutation t-test). When comparing DC microglia to sham, these values were 540% higher by “Cell” (g=3.0, p=0.0003 Bonferroni, p<0.0001 permutation t-test) or 433% by “Animal” (g=1.5, p=0.0251 Bonferroni; p=0.0540 permutation t-test). We also found evidence of a small increase in *trem2* mRNA expression in microglia contralateral to the injury compared to sham when compared by “Cell” (**Figure 3C**, p=0.0011 Dunns, 160% increase, g=0.9, p<0.0001 permutation t-test) although significance was not reached comparing “Animal” averages (p=0.8532, Bonferroni, 160% increase, g=1.34, permutation t-test p=0.0712) (**Data S1** in support of **Figures 3B-C**). Differences in total mRNA were mirrored by differences in *trem2* mRNA inside processes (**Figure 3E, Data S1**). In contrast, nuclear *trem2* content was unchanged among the four experimental groups (Cell: p=0.0501, Kruskal-Wallis; Animal: p=0.8716 one-way ANOVA) (**Figure 3D, Data S1**). The data suggest that higher TREM2 protein in the cytosol and membrane is due to higher expression of *trem2* mRNA in the microglial cell body and processes. Higher TREM2 protein in DC compared to On MN microglia contrasts with similar *trem*2 mRNA levels, suggesting post-transcriptional differences in *trem2* mRNA regulation and/or TREM2 protein half-life between DC and On MN microglia.

### TREM2 knock-out validation and suppression of downstream p-SYK

To test TREM2 roles in microglia interactions with MNs, we used two different mouse models to delete *trem2*: a global *trem2* Knock Out (GKO) and a timed and microglia-specific conditional *trem2* Knock Out (CKO). By comparing GKO and CKO we hoped to distinguish the effects of deleting *trem2* specifically in microglia just before PNI from any global effects of removing *trem2* from all tissues, and throughout development. TREM2 is expressed by microglia but also in many peripheral tissues including peripheral macrophages (Colonna, 2023; Jay et al., 2017; Lee et al., 2021; Sun et al., 2020; Wang et al., 2025). Moreover, TREM2 modulates several developmental actions, including synaptic refinement (Matteoli, 2024). To identify microglia in GKO animals, we crossed these animals with Tmem119^eGFP^ mice. To obtain animals with a microglia-specific deletion of *trem2,* we cross-bred mice with *trem2* floxed alleles with *tmem119-2A-creERT2* mice and the Ai9 R26 *lsl-tdTomato* cre-reporter mice. The triple transgenic (CKO) had all three alleles in homozygosis: *tmem119 ^creERT2/creERT2^*::*trem2^flx/flx^* ::*R26* ^lst-tdT/lsl-tdT^. Recombination was induced with five doses of tamoxifen (i.p. 2 mg/kg) separated by 24 hours, starting at P28. Nerve surgeries were performed at two months of age. From here on, Tmem119^eGFP^ animals with wildtype TREM2 will be referred to as wildtype.

TREM2 protein was undetectable in GKO and CKO mice microglia (**Figure 4A**; microglia genetic labels: eGFP in wildtype and GKO, tdTomato in CKO; both confirmed with IBA1). We further validated both models via RNAscope for *trem2* mRNA (**Figure 4B**). In this case microglia in GKO mice were identified with IBA1 antibodies only because eGFP signals with the Tmem119^eGFP^ reporter did not withstand RNAscope processing. This was not the case with tdTomato in CKO mice, which endured RNAscope and could be amplified with antibody labeling. 93.2% of IBA1+ cells express tdTomato in CKO mice suggesting high efficiency of tamoxifen-induced recombination in our model (N=4 animals, each analyzed in 5 L5 sections, **Figure 4C**). Moreover, the percentage of IBA1+ microglia in tdTomato CKO mice was similar to Tmem119^eGFP^ and CX3CR1^eGFP^ mice (**Figure 4C**). CKO mice showed no *trem2* mRNA signal in genetically defined microglia, while GKO mice showed a reduced signal in IBA1 microglia. This difference likely stems from differences in exon preservation based on the design of the two KO mouse models (see descriptions in methods). The RNAscope method was validated using positive and negative controls in serial sections from all samples (see methods). Thus, the lack of the *trem2* signal in GKO and CKO mice was not due to poor mRNA preservation in our samples or other technical inefficiencies.

To demonstrate reduced TREM2 microglia function in GKO and CKO mice, we analyzed the accumulation of downstream phosphorylated Spleen Tyrosine Kinase (p-SYK). p-SYK immunolabeling increased in parallel to the increase in TREM2 protein in activated wildtypeTmem119^eGFP^ microglia (**Figure 5A**). Since most p-SYK accumulated on the membrane, we outlined the microglia edge and measured p-SYK immunofluorescence integrated pixel density along the cell body membrane and normalized against background (see methods). Using this method, wildtype On MN microglia showed 62.5% increase in fluorescence and DC microglia a 737.4% increase compared to surveying microglia in sham controls (p<0.0001, Kruskal-Wallis followed by Dunn’s multiple comparisons; n=77 sham; n=104 On MN; n=24 DC; N=4 sham; N=6 cut) (**Figure 5B, Data S1**). Interestingly, contralateral microglia in PNI animals also showed a smaller (37%) but statistically significant, increase compared to sham (p=0.0028, Dunn’s test). This further suggests some small but functionally significant TREM2 upregulation in spinal microglia contralateral to the injury side. We also found that DC microglia in females had 125% more p-SYK than in males (p<0.0001, Dunn’s post-hoc comparisons; **Figure S4**). No sex differences in p-SYK accumulation were found in contralateral microglia or On MN microglia.

SYK phosphorylation occurs after TREM2-DAP12 activation, but numerous other receptors, including but not limited to C-type lectin 7a (CLEC7a), Fc Receptors, CD33, and CD22, also phosphorylate SYK in activated microglia (Ennerfelt et al., 2022; Pottorf et al., 2022; Wang et al., 2022). To corroborate that TREM2 loss resulted in reduced p-SYK signaling in ventral horn activated microglia, we compared GKO and CKO mice to our wildtypes (Tmem119^eGFP^ mice). Contralateral, On MN, and DC microglia had reduced p-SYK content in GKO and CKO mice compared to wildtypes. The results were confirmed in both “Cell” and “Animal” data sets (**Figure 5B, Data S1**). Contralateral microglia showed 52-53% reduction in GKO (p<0.0001 Cell; p=0.0061 Animal) and 35-38% in CKO (p<0.0001 Cell; p=0.0335 Animal). On MN microglia showed 58-58% reduction in GKO (p<0.0001 Cell; p=0.001 Animal) and 49-48% in CKO (p<0.0001 Cell; p=0.0335 Animal). Finally, DC microglia showed the largest reduction; 88% in GKO (p<0.0001 Cell) and 87% in CKO (p<0.0001 Cell).

In summary, TREM2 is efficiently removed from microglia in both GKO and CKO mice, and this results in lower p-SYK accumulation in activated microglia after PNI, particularly in DC microglia. Within these large reductions in p-SYK, female microglia retained more p-SYK and this reached significance in CKO (p=0.002, Dunn’s post-hoc comparisons; **Figure S4**), suggesting parallel pathways generating p-SYK are more active in female microglia.

### TREM2 knockout does not affect the number, territories, or process lengths and branching of non-activated surveying microglia in sham animals

TREM2 KO was previously reported to alter surveying microglial morphology, reduce the number of microglia, and collapse microglia territories in one-month-old murine cortex (Jay et al., 2019). We sought to confirm these findings in TREM2 GKO and CKO ventral horn spinal cord microglia compared to Tmem119^eGFP^ wildtype mice (N=7 Tmem119^eGFP^ wildtype; N=7 Tmem119^eGFP^ GKO; N=5 Tmem119^creERT2^ tdTomato CKO). No sex differences were found for the number of microglia or average microglia territory size in the L5 ventral horn of uninjured mice (wildtype N=3 male/4 female; GKO N=4 male/3 female; CKO N=4 male/1 female; **Figure S5**). Estimated microglia numbers were on average 5.3 ±0.8 (±SD) per 10^6^ µm^3^ (a 100 µm side cube volume), and their 2-dimensional projection territories in wildtypes measured on average 4,007 µm^2^ ±707 (SD), corresponding to approximate diameters of 65-77 µm, suggesting that processes extending further than 50 µm from the cell body partially overlap. Although there was a slight trend (∼11-12% change) towards reduced microglia numbers (**Figure 6A-B**) and increased average territory sizes (**Figure 6C**) in the ventral horn of GKO and CKO sham mice compared to wildtype, these differences were not significant (one-way ANOVAs p=0.2246 microglia number, p=0.3871 microglia territories; **Data S1** in support of **Figures 6A and 6B**). This conclusion was confirmed with estimation statistics (**Data S1** in support of Figures 6A and 6B). Cell morphology did not change either in GKO sham animals (Sholl analyses of n=5 microglia reconstructed and skeletonized compared to wildtype microglia reconstructions from **Figure 1**, N=6 per condition, equal sex; **Figures 7A,B**). Two-way ANOVA for the number of intersections at different Sholl distance bins in different genotypes found significant differences with Sholl bin (p<0.0001, as expected) but not according to genotype (p=0.6677) or the interaction of Sholl bin and genotype (p=0.0571) (n=30 reconstructed cells in GKO and WTs, **Data S1** in support of **Figure 7B**). We could not adequately compare CKO microglia because tdTomato labeling filled distal processes less efficiently than eGFP

In conclusion, TREM2 deletion had no discernible effects on the morphological properties of ventral horn microglia before activation.

### TREM2 knockout does not affect microglia proliferation following nerve injuries

TREM2 KO has been shown to reduce microglia proliferation and migration in Alzheimer’s disease (Wang et al., 2020; Zheng et al., 2017) and cuprizone-induced demyelination models (Cantoni et al., 2015). Therefore, we performed a time-course analysis of ventral horn microglia proliferation by estimating their numbers (**Figure 6D,E**) and the percentage of microglia expressing Ki-67 (**Figure 6F,G**) after PNI in both TREM2 KO models (GKO or CKO). Wildtype, GKO, and CKO mice underwent sciatic nerve cut repair with a Fast Blue nerve soak to retrogradely label the entire injured sciatic MN population. Mice were euthanized at 3, 7, 14, 21, and 28 dpi (N=4 animals per genotype/time point, 2 male/2 female, except for CKO animals at 21 dpi in which only 2 females were analyzed). Sham mice were not included in these analyses as Fast Blue nerve soak labeling requires nerve transection.

The number of proliferating microglia in the ventral horn after PNI (estimated by quantifying n=5 L5 sections per animal) showed significant differences analyzed for time after injury and genotype (WT, GKO, and CKO) (two-way ANOVA; time after injury p<0.0001; genotype p=0.0157) (**Figure 6E**, **Data S1)**. In all three genotypes, the number of microglia peaked at 14 dpi and then declined towards normal values by 28 dpi similar to our previous reports in different animal cohorts with different genetic labels (Rotterman et al., 2024). Although a genotype effect was detected and GKO and CKO mice consistently trended towards fewer microglia than wildtype mice, post-hoc pairwise Bonferroni tests did not detect significant differences between genotypes at any time point (**Data S1** in support of **Figure 6E**). The low n per time/point genotype combined with inter-animal variability suggested underpowered comparisons given the relatively small effect sizes (always smaller than 21% difference during the rising phase of microgliosis from 3 to 14 dpi). Power analyses (G*Power version 3.1.9.7) indicated that n’s would need to be increased to 40 animals per genotype/time point, which is unfeasible. The differences among genotypes are therefore small, if any. The percentage increase in cell numbers from 3 to 14 dpi was similar in all three genotypes (wildtype 68%; GKO 66%; CKO 56%). We conclude that TREM2 KO had only a small effect on microglia proliferation in the spinal cord after PNI in contrast to other pathologies/injuries in which TREM2 was reported to more strongly reduce microglia proliferation (Cantoni et al., 2015; Wang et al., 2020; Zheng et al., 2017).

We confirmed this conclusion using Ki-67 antibodies as a direct marker of cell-cycling microglia. The Ki-67 antibody used was made in mouse and cross-reacted with microglial Fc receptors upregulated after activation. Despite this cross-reaction, we were able to distinguish between specific and nonspecific Ki-67 labeling by focusing on true nuclear localization. In all genotypes, the proportion of microglia expressing Ki-67 was highest at 3 dpi (25.4% ±3.0, 33.0% ±17.4, and 15.8% ±8.8 (±SD) in WT, GKO, and CKO, respectively) and much less at 7 dpi (3% ±0.5, 3.7% ±2.9, and 0.9% ±0.9) (**Figure 6F,G**). Less than 1% of microglial cells showed Ki-67 at later time points. Two-way ANOVA showed significant differences for time after injury (p<0.0001) but not among genotypes (p=0.1156) (**Data S1** in support of **Figure 6G**). This result agrees with our previous data showing that ventral horn microglia proliferation is restricted to the first week after injury, being higher at 3 dpi compared to 7 dpi (Rotterman et al., 2019). Interestingly, remaining Ki-67+ microglia at 7 dpi seem to depend on physically interacting with axotomized MNs. Around a third of proliferating Ki-67+ microglia were found On MN at 3 dpi, but this percentage increased to 75.3% ±29.0 at 7 dpi. Death clusters rarely contained any proliferating Ki-67+ microglia in any animal or genotype, suggesting that death clusters form by coalescence of On MN microglia when the MN degenerates, and not by local proliferation.

We next investigated whether TREM2 is necessary for the formation of death clusters. The number of death clusters at 3, 7, 14, 21, and 28 dpi was compared in wildtype versus GKO and CKO. In each spinal cord and animal, we counted death clusters in all lumbar sections showing microgliosis. This number was normalized to the number of sections collected and multiplied by 100 to estimate the total number of death clusters in a 5 mm length (100 x 50 mm thick sections) of the lumbosacral enlargement. This region corresponds to the column of injured sciatic MNs, which encompasses Lumbar segments 4 to 6. We did not detect sex differences (**Figure S6**) and therefore pooled data from both sexes. In wildtype mice (Tmem119^eGFP^), the number of death clusters peaked at 14 dpi (**Figure 6H-I**), coinciding with the peak of microgliosis. Although GKO and CKO tended to have fewer death clusters than wildtype mice, the differences between wildtype, GKO, and CKO were not significant at any time point (two-way ANOVA for time after injury and genotype: p<0.0001 time after injury; p=0.3989 genotype; **Figure 6I** and **Data S1** (in support of **Figure 6H**) show comparisons among genotypes at each time point). The result was confirmed with estimation statistics at 14dpi (wildtype vs GKO Hedges’ g =-1.2, permutation t-test p=0.1176; wildtype vs CKO Hedges’ g=0.1, permutation t-test p=0.801) (**Figure 6H**). We conclude that the formation of death clusters is largely unaffected by TREM2 deletion.

### TREM2 KO has only limited effects on microglia morphology and the number of microglia interacting with injured MNs

Next, we analyzed activated microglia morphology, migration, and interactions with MNs 14 days after sciatic nerve-cut ligations in the absence of TREM2 (**Figure 7A-B**, n=30 cells in wildtype and 30 in GKO for each microglia category, 5 cells per animal N=6 wildtypes and 6 GKO animals, equal sexes). GKO microglial morphology, as assessed by Sholl intersection analysis, did not differ from wildtype microglia in sham, contralateral, or “On MN” microglia (two-way ANOVAs with factors Sholl distance and genotype: genotype p = 0.6677, 0.1016, and 0.3120, respectively; Sholl distance was highly significant in all cases, p < 0.0001) (**Figure 7B**; **Data S1**). Pairwise Bonferroni post hoc tests revealed no significant differences between wildtype and GKO mice at any distance point across these three microglial populations, with the exception of the 20 µm radius in contralateral microglia (p = 0.029). However, GKO DC microglia showed significant differences compared to wildtype (two-way ANOVA; genotype p=0.0014, Sholl bin p<0.0001). Specifically, there was a 34%, 83% and 105% increase in intersections, respectively at 10, 20, and 30 µm Sholl distances, but only the increase at 20 µm reached significance using Bonferroni pair-wise comparisons (p=0.0187) (**Figure 7B, Data S1**). We conclude that TREM2 was minimally involved in the morphological changes that occur in On MN microglia although TREM2 absence in GKO partially prevents process retraction in DC microglia. Analyses were not performed in CKO microglia because as explained before, tdTomato and eGFP labeling were not comparable in the extent to which they fill distal processes.

Microglia migration and microglia-neuron interactions have both been shown to depend on TREM2 activity in other neuropathologies (Jairaman et al., 2022; Mazaheri et al., 2017; Sayed et al., 2018) and therefore we examined in all three genotypes the number of microglia in contact with injured and uninjured MNs. In sham control animals, most contacts with MNs occurred via microglia processes, while after PNI, microglia and MN cell bodies frequently apposed to each other in either the On MN or DC configuration. Analyses were performed on Fast Blue MNs retrogradely labeled from the LG muscle and contacted by either eGFP (wildtype and GKO) or tdTomato (CKO) labeled microglia. We analyzed 8 wildtype (equal sex), 7 GKO (4 male /3 female), and 5 CKO (2 male /3 female) animals with sham surgeries and 8 wildtype (equal sex), 8 GKO (equal sex), and 7 CKO (4 male /3 female) animals after PNI. We only detected a sex difference in the number of microglia forming part of DC clusters. Male DC microglia contain on average 5.4 ± 1.7 microglia (N=4 animals) while female microglia had 7.6 ±0.6 microglia (N=3 animals), a 42% significant increase (p= 0.0374, Bonferroni post-hoc test, **Figure S7**). Because of small N and the fact that no significant differences were found in other microglia phenotypes we pooled data from males and females together. After sham surgeries, the number of microglia contacting MNs was 1.6 ±0.6 (±SD) in wildtype, 1.7 ±0.7 in GKO, and 1.1 ±0.5 in CKO, but after PNI, this number increased to 5.3 ±2.4, 5.7 ±0.9, 4.4 ±0.7 microglia On MN and 6.3 ±1.7, 6.7 ±1.8, 6.6 ±2.0 in DC microglia. The number of MNs analyzed per animal (excluding outliers) was 7 to 10 for sham or On MN microglia (9.7 ±1.0 across all three phenotypes). However, the number of death clusters found and analyzed per animal was less (1 to 10, for an average of 3.9 ±1.0). Thus, the total number of MNs analyzed in each group was 75 wildtype, 69 GKO, and 58 CKO in animals with a sham surgery, and 77 wildtype, 66 GKO, and 78 CKO MNs after PNI (On MN phenotype), and 33 wildtype, 22 GKO, and 29 CKO death clusters. Data was analyzed in “Cell” and “Animal” data sets. Average numbers of microglia associated with each MN (sham or On MN) or in DC microglia in “Cell” data were similar to those calculated for “Animal” averages, but as expected, there was higher variance (**Figures 7C,D**). We independently compared the number of microglia associations in each experimental condition (sham, On MN or DC) across genotypes (**Figure 7C**) and the microglia associations in sham, On MN, and DC microglia for each genotype (**Figure 7D**; full statistical data in **Data S1)**. The number of microglia associations with MNs after PNI, comparing On MN and DC microglia to the sham condition, was increased in all three genotypes in both “Animal” and “Cell” data sets (p<0.0001 in all cases; one-way ANOVAs followed by pair-wise Bonferroni tests for “Animal” data, Kruskal-Wallis followed by Dunns tests for “Cell” data). This conclusion was confirmed with estimation statistics resulting in large g values from 3.3 to 5.3 in “Animal” data and 2.6 to 3.7 in “Cell” data. We also detected a small decrease in associated microglia in the CKO genotype compared to wildtype and GKO in both sham and On MN experimental conditions. This difference reached significance in the “Cell” data set but not in the “Animal” data. Estimation statistics confirmed the significance obtained with “Cell” data but also showed that effect sizes were small (g varied from 0.44 to 0.63). Given the small effect and the fact that the GKO condition did not replicate CKO results, we conclude that this small difference is most likely derived from the fewer distal processes revealed with tdTomato genetic labeling in CKO animals. Accordingly, no difference in the number of microglia associations was observed across genotypes in death clusters in which the efficiency of process labeling is irrelevant.

In conclusion, TREM2 KO (global or microglia-specific) had only small effects on the morphological changes that occur in microglia from surveying to activated On MNs or DC microglia, and no significant effects were detected on microglia migration and association with injured MNs. Taken together with previous results, we conclude that TREM2 is not necessary for microglia proliferation, migration towards or enwrapping injured MNs, or the evolution of microglia interactions from On MN phenotypes towards death cluster formation.

### Death Clusters display the greatest CD68 expression regulated by TREM2, especially in females

DC microglia are morphologically similar to highly phagocytic Disease-Associated Microglia (DAM) with high expression of *trem2* (Deczkowska et al., 2018; Keren-Shaul et al., 2017; Sasaki et al., 1997; Sobue et al., 2021; Streit et al., 1999). Thus, we tested whether DC microglia show a similarly strong phagocytic phenotype and whether this is regulated by TREM2. We used CD68 expression to monitor phagocytic phenotypes and analyzed microglia at the peak of their activation (14 days after a sciatic cut-ligation). Like experiments on microglia morphology, we only compared wildtypes to GKO mice because of differences in microglia labeling in CKO mice due to differences in genetic labeling properties (Tmem119^tdTomato^ in CKO vs Tmem119^eGFP^ in GKO and WTs). EGFP vs tdTomato labeling can bias results because they result in different microglia volume estimates.

As expected, microglia activated after PNI increased CD68 expression, with DC microglia showing the highest CD68 levels (**Figure 8A).** TREM2 absence (GKO) reduced CD68 content, but important sex differences were noted. To quantify this result, we analyzed 8 wildtypes (4 male / 4 female) and 7 GKOs (4 male / 3 female) in sham surgery animals and 7 wildtypes (4 males / 3 female) and 8 GKOs (4 male / 4 females) 14 days after PNI (sciatic cut-ligation). We analyzed between 17 to 26 microglia in each animal and at each of the four locations (sham, contralateral, On MN and DC) for a total of 464 IMARIS 3D reconstructed microglia in wildtypes and 488 in GKO animals. Sample sizes for each experimental condition after identifying outliers are reported in **Data S1** in support of **Figures 8 and 9** data analyses. We calculated the proportion of microglia volume occupied by CD68 granules (**Figure 8B**, **Figure 9A,D**), the number of CD68 granules per microglia (**Figure 8C**, **Figure 9B,E**), and the maximum size of detected CD68 granules in each cell (**Figure 8D**, **Figure 9C,F).** We used maximum size instead of average size because microglia contained mostly small granules, sometimes below optical resolution limits (<0.2 µm), and these large populations of small-sized granules overwhelmed average size estimations. As before, we generated “Cell” and “Animal” data, but for simplicity, we report statistical tests on “Cell” data. We noted significant sex differences in wildtypes (**Figure 8B,C,D; Data S1** in support of **Figure 8B**) and therefore analyses of CD68 expression in the absence of TREM2 were performed in males and females separately (**Figure 9**).

“Cell” data two-way ANOVA for microglia type and sex showed significance for both (p<0.0001) in total volume and number of granules in sham, contralateral, and On MN microglia, but not in DC microglia. Pairwise comparisons showed that females had more CD68 content in surveying (sham and contralateral) and activated (On MN) microglia compared to males (usually p<0.0001 Bonferroni tests confirmed by estimation statistics, **Data S1** in support of **Figure 8B**). Sex differences in CD68 content were not extensive in DC microglia. The percentage of microglia cell volume occupied by CD68 granules in wildtype female sham, contralateral, On MN and DC microglia was respectively 10.0% ±7.9 (SD), 8.4% ±5.3, 9.4% ±5.1, and 37.5% ±20.0 and in wildtype male microglia 3.2% ±2.0 (SD), 3.8% ±3.2, 6.7% ±7.3, and 29.2% ±23.2.CD68. Thus, female and male DC microglia had two-to-three times more CD68 granules and total CD68 volume compared to the same sex surveying and activated On MN microglia. However, overall, female microglia had more CD68 granules and total CD68 volume than males. In contrast, granule maximum size did not show consistent differences according to sex in sham, contralateral, and On MN microglia (Bonferroni tests), although they reached significance when compared using estimation statistics and permutation t-tests (**Data S1** in support of **Figure 8B).** Nevertheless, estimated differences in average granule maximum size were small and highly variable from animal to animal (female to male Hedges’s g=1.2 in sham, 0.8 contralateral, 0.4 On MN microglia).

To test the role of TREM2, we compared CD68 content in wildtype and GKO animals in all four microglia types (sham, contralateral On MN and DC) and separated the data according to sex (**Figure 9**). Two-way ANOVAs in wildtype “Cell” data on total volume, granule numbers and maximum granule sizes detected significant changes according to animal genotype, microglia phenotype, and their interaction in both sexes (**Data S1** in support of **Figure 9**). “Animal” data statistics generally agreed (**Data S1**). DC microglia maximally expressed CD68 in both wildtypes and GKO mice. In wildtypes, DC microglia showed CD68 granule volume increased by 286-326% (males) and 925-985% (females) compared to sham or contralateral microglia, respectively (p<0.0001 in all comparisons, Bonferroni t-tests; **Figure 9A,D**; **Data S1**). These differences were driven by a large increase in the average maximum size of CD68 granules (usually over 1,000% compared to sham or contralateral microglia, p<0.0001 in all comparisons; **Figure 9C,F, Data S1**) while changes in CD68 granule numbers were less intense and frequently did not reach significance (**Figure 9B,E, Data S1**). Differences between sham and contralateral and On MN microglia for all three estimates were smaller, more variable, or non-significant (**Data S1** in support of **Figure 9** statistical analyses). Thus, CD68 granules in DC microglia are much larger in both sexes, confirming a different phagocytic behavior. In addition, the difference is larger in female compared to male DC microglia. TREM2 deletion had larger effects in females (**Figure 9A-C**) compared to males (**Figure 9D-F**). In females, decreases in total CD68 volume occupancy compared to wildtypes were relatively similar in effect size in sham (76.3% reduction, g=-1.4), contralateral (69%, g=-1.6), On MN (47.1%, g=-1.0) and DC microglia (69.1%, g=-2.3). Decreases were always significant (p<0.0001 Bonferroni tests, and similar p values were obtained with permutation t-tests, except for the difference in DC microglia p=0.0004, **Figure 9A, Data S1**). These decreases were paralleled by reductions in granule number and average maximum sizes (details in **Data S1** in support of **Figure 9**). In females, decreases in granule number (70.6%, g=-2.1) and average maximum size (86.2%, g=-2.2) were consistently higher in DC microglia compared to the other microglia types (sham, contralateral and On MN). Thus, the Hedge’s g estimate for CD68 reductions in female TREM2 GKO was larger for all CD68 content parameters measured in DC microglia compared to other microglia types.

The results in males were more complex. Total CD68 volume was reduced in TREM2 GKO DC male microglia (45.7%, g=-0.7), but results were inconsistent between Bonferroni tests (p<0.0001) and permutation t-tests (p=0.1284). Parallel reductions in CD68 granule numbers (52.0% g=-1.7) and maximum size (21.2%, g=-0.2) in male TREM2 GKO DC microglia were found, but they were generally non-significant by both Bonferroni and permutation t-tests. Surprisingly, we detected consistent increases in CD68 granule number in sham (86.2%, g=1.0), contralateral (97.1%, g=1.0), and On MN (76.3%, g=1.0) microglia of TREM2 GKO males. These differences were always significant compared to wildtype controls (p<0.001 for all comparisons with both Bonferroni and permutation t-tests). A change in granule maximum size in the opposite direction did not reach significance but seemed to counteract granule numbers, such that overall, no change was found in male microglia volume occupancy by CD68 granules.

Collectively, the data show that DC microglia are phagocytic hubs and CD68 granules appear dysregulated in both female and male TREM2 GKO mice. But while the lack of TREM2 reduced CD68 in all microglia types in females, and more intensely in DC microglia, the effects were different in males. Male GKO microglia, other than in death clusters, consistently showed more granules but without significant changes in total volume, suggesting the excess CD68 granules were generally smaller. Male DC microglia were not significantly affected by the lack of TREM2, suggesting that alternative signaling pathways support CD68 expression in males. Larger TREM2 GKO effects in female DC microglia parallel their larger expression of *trem2* mRNA and TREM2 function (p-SYK accumulations) shown before.

### Microglia TREM2 is necessary for normal motoneuron cell body swelling during the chromatolytic reaction following axotomy

Healthy mouse MNs exhibit cell size bimodal distributions corresponding to small γ-MNs and large α-MNs, that can also be distinguished by NeuN Neuronal Nuclei (NeuN)-immunoreactivity (Shneider et al., 2009): low in γ-MNs and high in α-MNs. We confirmed this distribution in wildtype naïve mice lumbar 5 sections immunostained with Choline Acetyltransferase (ChAT) and NeuN antibodies (N=3; 1 male, 2 females). ChAT+ MNs were imaged, classified as low or high NeuN, and their cell body sizes measured (**Figure 10A,B, Data S1**). Five sections were analyzed per animal, averaging 145.3 ±3.2 (SD) MNs per animal. On average, 33.5% ±7.9% were low-NeuN MNs. Cell size distributions were obtained by binning area estimates (see methods) in 50 µm^2^ increments for all MNs and, independently, for low and high NeuN MNs (**Figure 10B**). Cell size distributions were fitted to Gaussian curves, pooling together all MNs sampled from the three animals (n=436 MNs, 290 high-NeuN, and 146 low-NeuN). Optimal correlations were obtained with bimodal models (r^2^=0.8280 for all-MNs, 0.8377 for low-NeuN MNs, and 0.8286 for high-NeuN MNs). The low- and high-size peaks in the all-MN size distribution correlated with the low-NeuN and high-NeuN populations, reflecting the γ- and α-MN populations (dashed lines in **Figure 10B**). The γ-MN peak was estimated at a cross-sectional area of 289.8 µm^2^ ±8.4 (SEM). The average area independently estimated for low-NeuN MNs in the three animals was higher (348.4 ±34.1, N=3), and this difference can be best explained by a population of low-NeuN MNs that skew towards larger sizes. The estimated cross-sectional area of the α-MN peak was 738.0 µm^2^ ±14.7, which was no different from the average value of all high-NeuN MNs in the three animals (797.6 ±42.7, ±SEM, N=3). These data were compared to the cell sizes of Fast Blue-labeled sciatic MNs at different times after nerve injury and axotomy.

At 3, 7, 14, 21 and 28 days post-injury (dpi) we analyzed wildtype, GKO, and CKO animals with genetically labeled microglia after nerve injuries and Fast Blue labeling of axotomized MNs (**Figure 10C**) (N=4, 2 male / 2 female at all time points and genotypes, except for GKO at 28 dpi in which N=3, 1 male and 2 females, and CKO at 21 dpi N=2 females, total N=57 mice). There was no sham control group because a nerve cut is necessary to retrogradely label MNs with a Fast Blue soak of the nerve stump. NeuN-immunoreactivity was not tested because axotomy downregulates NeuN in MNs, and it is not recovered until MNs reinnervate muscles (Alvarez et al., 2011). Cell-size distributions were analyzed for the presence of bimodal peaks (**Figure 10E**). Thereafter, the data were transformed into cumulative plots to statistically test differences using Kolmogorov-Smirnov (K-S) tests at different post-injury times (**Figure 10F**). In all animals, the cell size distributions differed from normal in that the double peak distribution was smeared, suggesting changes in size in all types of axotomized MNs, as early as 3dpi. In wildtypes, we noted a reduction of the lower-size peak (almost disappearing at 14 dpi) and a shift of the whole distribution towards larger sizes from 3 to 14 dpi, and then a return at 28 days towards the 3-day distribution (**Figure 10E**). Thus, estimates of average sizes increased from 3 to 14 dpi by 17.3% in the low-size peak and 24.4% in the high-size peak, returning towards the 3-day value at 28 days (**Data S1** in support of **Figure 10E)**. These changes in cell size were blunted in GKO and CKO animals (**Figure 10E**, **Data S1**). Correspondingly, K-S tests indicated that cell size distributions at 7 and 14 days were significantly different from all others in wildtypes, but not in GKO or CKO animals (**Figure 10F**; **Data S1**).

Examination of the size distributions suggested that a major change occurred in cells larger than 1,000 µm^2^ in cell area, corresponding to cell swelling in the α-MN group. Thus, we calculated the percentage of MNs with areas larger than 1,000 µm^2^ in all animals and tested for significant differences according to time after injury (dpi) and genotype (**Figure 10D**). A two-way ANOVA for both variables returned statistical differences for both (p<0.0001), and post-hoc Bonferroni tests found statistically significant differences from 3 dpi to 7 and 14 dpi in wildtypes returning towards baseline at 21 and 28 dpi (**Data S1** in support of **Figure 10D**). In contrast, changes in cell size were smaller in GKO and CKO, and significant differences were only detected between 3 and 28 dpi compared to 14 dpi (**Figure 10D, Data S1**). We confirmed the smaller change in size in the knock-out animals through Hedge’s g estimates; comparison between 3 and 14 dpi was large in wildtypes (g=4.4) and small in GKO (g=0.9) and CKO (g=2.1). Consequently, significant differences occurred when comparing cell size at 14 dpi wildtypes to GKO and wildtypes to CKO (p<0.0001 in both cases), while there were no significant differences between GKO and CKO mice (p>0.9999) (**Figure 10D** and **Data S1**).

In summary, the microglial reaction coincides in time course with transient changes in MN cell size following axotomy that are believed to be reflective of a healthy regenerative response with exponential increases in RNA and protein synthesis and metabolism (Fu & Gordon, 1997; Lieberman, 1971). Microglial TREM2 seem necessary for normal changes in MN cell size to occur after axotomy, suggesting a possible role in facilitating regeneration.

### Microglia TREM2 activity may remove a proportion of small motoneurons

To estimate the distribution of α- and γ-MNs after 8 weeks of peripheral nerve regeneration we further analyzed wildtype animals (Tmem119^eGFP^) after a sham (N=6 equal sex number, n=1,286 MNs) or a cut-repair nerve injury (N=3 all-female, n=464 MNs) and compared their cell size distributions and NeuN expression to the injured and uninjured sides of CKO animals (N=2 animals 1 male and 1 female, n=497 MNs in injured side and n=459 MNs in the uninjured side) after a similar regeneration period (**Figure 11**). Sham control animals and the uninjured side of experimental animals demonstrated α-MN and γ-MN peaks similar to naïve animals (**Figure 10B**), and these correspond respectively, to MNs expressing high or low NeuN. However, 8 weeks after injury the animals consistently showed a partially reduced γ-MN peak in the side ipsilateral to the injury and a spread of NeuN negative cells towards the α-MN distribution (**Figure 11A**). Both effects were abolished in two CKO animals (one male and one female) that showed 8 weeks after the nerve injury normal α-MN and γ-MN peaks and NeuN high and low MN distributions (**Figure 11B**). Cell size distribution fitting demonstrated that in injured wildtypes the smaller size peak contains fewer cells and shifts towards larger cell sizes (**Figure 11C**). This is best interpreted as the loss of some cells within the original γ-MN peak and the appearance of a population of MNs with cell sizes intermediate between α- and γ-MNs. In contrast, the γ-MN peak is in CKO animals is preserved and MN size distributions in CKO animals perfectly overlap between the injured and uninjured side of the spinal cord (**Figure 11D**). These results suggest a partial loss or changes in cell sizes of subpopulations of γ-MN cell bodies after nerve transection injury that are rescued by selectively deleting *trem2* from microglia with results similar in male and female mice.

### TREM2 deficiency slows muscle reinnervation by α-motor axons

To confirm a role for TREM2 modulating α-MN regeneration after axotomy, we analyzed muscle reinnervation at eight weeks post-sciatic cut-repair, a time point at which most neuromuscular junctions (NMJ) are expected to be re-innervated in this nerve injury model (Rotterman et al., 2024). To assess muscle innervation, we performed immunohistochemistry for Neurofilament Heavy Chain (NFH) and Vesicular Acetylcholine Transporter (VAChT) on alpha-Bungarotoxin labeled NMJs (α-BTx) of the ipsilateral LG muscle (**Figure 12A**). Then we quantified coverage by VAChT (motor axon end-plate) of the NMJ area defined by α-BTx (see methods). NFH immunostaining was useful to qualitatively evaluate preterminal axons. VAChT signals were weaker and more variable than α-BTx and based on control data we established that NMJs with over 50% VAChT coverage of the α-BTx signal are fully innervated. Using this criterium we estimated the percentage of fully innervated NMJs by sampling 50 NMJs per animal. No sex differences were identified. Wildtype and GKO “controls” consisted of sham mice, whereas CKO “controls” were a mix of two sham and one naïve animal to increase the N for statistical analysis (N=6 wildtype, 4 GKO, 3 CKO for controls; N=5 wildtype, 6 GKO and 4 CKO mice with cut and repair injuries). Two-way ANOVA detected significant differences in NMJ innervation according to genotype (p=0.0020), time after injury (p=0.0017) and their interaction (p=0.0076) with the percentage of reinnervated NMJs being significantly lower in GKO and CKO mice compared to wildtypes (**Figure 12B**). Compared to regenerating wildtypes, GKO muscles had 37% less fully innervated NMJs (p=0.002, g=-2.19), and CKO muscles had 56% less (p=0.0002, g=-3.68) (**Figure 12B, Data S1**). No significant difference was found between GKO and CKO for NMJ innervation (p=0.4798 Bonferroni’s g=-0.75). Similarly, no significant difference in NMJ innervation was found in wildtypes between control and injured mice (p=0.6630, Bonferroni tests) (**Figure 12C, Data S1**), suggesting full reinnervation at this time point. In contrast, GKO cut-repair LG muscles (N=6) had 23% less fully innervated NMJs compared to GKO controls (p=0.0525 Bonferroni test, g = -0.97 permutation t-test p=0.127) (**Figure 12C, Data S1**), and CKO cut-repair LG muscles had 54% less fully innervated NMJs compared to CKO controls (p=0.0005 Bonferroni test, g=-2.97, permutation t-test p<0.0001) (**Figure 12C, Data S1**). The results, therefore, suggest that motor axon regeneration and NMJ reinnervation are delayed in the absence of TREM2.

## Discussion

The significance of microglia enwrapping the cell bodies of MNs axotomized after nerve injuries, and whether microglia interactions favor regeneration or damage the MNs, is unclear. Our results suggest microglia exert dual functions over different axotomized MNs, both regulated by TREM2. More frequently, microglia surround MN cell bodies while remaining individualized, and these MNs show cell body changes suggesting a normal reaction associated with regeneration. Other microglia form tight clusters surrounding degenerating MNs. TREM2 is expressed in microglia involved in both types of interactions, although it is more highly expressed in “death cluster” microglia. To investigate TREM2 function we used two knockout models: a global knockout and a microglia-specific TREM2 deletion induced before nerve injuries. TREM2 deletion in both models did not affect microglia-activated morphologies, proliferation, migration towards MNs, number of interactions with MNs, or formation of microglia clusters surrounding dying MNs. TREM2 deletion did however reduced CD68 expression of microglia in death clusters and preserved the small sized MN peak of putatively γ-MNs. Moreover, TREM2 deletion prevented cell body swelling characteristic of α-MNs undergoing regeneration-associated metabolic changes, and this correlated with delays in muscle reinnervation. The results suggest that TREM2 function is distinct in each of the two microglia phenotypes: it promotes regenerative mechanisms over regenerating MNs, while it facilitates phagocytosis of degenerating MNs.

### TREM2 deficiency prevents motoneuron cell body swelling after axotomy and slows regeneration

Whether microglia facilitate the regeneration of axotomized MNs and their motor axons has been controversial. One study noted that microglia surrounding axotomized MNs are enriched in MHC-II (Major Histocompatibility Complex 2) and proposed they promote regeneration by the capacity of MHC-II expressing microglia for releasing cytokines and growth factors (Gehrmann et al., 1991). This agrees with a study that found slower motor axon growth when the microglia reaction around axotomized MNs is diminished after IL-6 (Interleukin 6) deletion (Galiano et al., 2001). A more recent study suggested that microglia release Ciliary Neurotrophic Factor (CNTF) over axotomized MNs to promote their regeneration (Tanaka et al., 2017). Intriguingly, when CNS neurons (that regenerate poorly) were grafted into peripheral nerves their cell bodies became surrounded by microglia and axon growth was enhanced (Shokouhi et al., 2010). In contrast, opposite conclusions were reached in studies that prevented microglia proliferation around axotomized MNs by interfering with CSF1 or inhibiting microglia proliferation (Horvat et al., 2001; Kalla et al., 2001; Svensson & Aldskogius, 1993b).

Our results suggest that microglial TREM2 is necessary for MN cell body swelling after injury, and this correlates with slower muscle reinnervation. Cell body swelling is central to the retrograde chromatolytic reaction associated with increased gene transcription, RNA metabolism, and protein synthesis (Fu & Gordon, 1997; Lieberman, 1971). Cellular changes include reorganization of the Nissl substance, migration of the nucleus to the cell periphery, and enlargement of the nucleolus, all associated with cell body swelling. In mice, the chromatolytic reaction is not as clear as in other species (Lieberman, 1971), but cell swelling remains a prominent change informing about metabolic modifications associated with regeneration. Lack of TREM2 impaired this reaction, suggesting a regenerative deficit confirmed by the slower re-innervation of muscle NMJs. Similar results in TREM2 global and microglia cell-specific knockouts rule out explanations based on the function of TREM2 regulating Schwann cell pro-regenerative phenotypes (Zhang et al., 2024).

Microglia may modulate MN regeneration by their known roles in monitoring and modulating neuron bioenergetics. In the adult murine cortex, surveying microglia sample the state of neuronal mitochondria through specialized microglia-neuron junctions formed between Kv2.1 clusters on neuronal membranes and P2Y12 receptor clusters in microglia (Cserép et al., 2020). Preventing their formation hampers neuronal mitochondrial function. Moreover, after acute injury the number of these microglia contacts increases and exert neuroprotection. Similarly, microglial TREM2 deficiency causes a deficit in neuronal bioenergetics, abnormal mitochondrial ultrastructure, and delayed maturation in CA1 pyramidal neurons (Tagliatti et al., 2024).

We also noted that microglia surrounding MNs can transition from “On MN” phenotypes to death cluster microglia, and in this case, TREM2 promotes phagocytosis. TREM2 and microglia functions might then be a continuum from pro-regenerative to pro-phagocytic depending on the health status signaled by the MN. These signals are currently unknown, but one possibility is the many neuropeptides upregulated in MNs after axotomy. For example, pituitary adenylyl-cyclase-activating peptide (PACA) is upregulated in axotomized MNs and modulates microglia function such that mice lacking PACA have an exaggerated microglia reaction impairing axon regeneration and increasing MN loss (Armstrong et al., 2008).

### TREM2 deficiency decreased CD68 in microglia, particularly in death clusters and preserved small size motoneurons

Microglia tight cell clusters aggregating over spinal MNs showing morphological features of degeneration attract CD8 cytotoxic T-cells (Rotterman & Alvarez, 2020) and are similar to microglia and T-cells associations reported in the facial nucleus after nerve transections that also induce mixed MN fates; some undergo regeneration while others degenerate (Ha et al., 2006; Ha et al., 2008; Kalla et al., 2001; Raivich et al., 1998). In the facial nucleus, these aggregations were named “microglia nodules”, and a similar term is used for clusters of microglia and CD8 T-cells associated with neuronal death during encephalitis or virus infections (Laukoter et al., 2017; Troscher et al., 2019) or with axon de-myelination and die-back in multiple sclerosis (van den Bosch et al., 2024). Microglia nodules express cytokines that attract T-cells (*Ccl5, Ccl2, Ccl7, Cxcl10, Cxcl11 and Cxcl16*), and after T-cell recruitment microglia properties are modified by T-cell interferon-γ (Troscher et al., 2019). Our data show they become enriched in TREM2 and CD68 content but with important sex differences: both are higher in females. CD68 content was reduced in TREM2 knockouts, with the effect being also larger in females. This suggests that death cluster microglia in females heavily rely on TREM2 to upregulate phagocytic activity, while similar microglia in males may use alternative or parallel signaling mechanisms. Similarly, RNA sequencing data indicated that in a mouse sciatic nerve chronic constriction injury model, dorsal horn female microglia utilize TREM2 signaling while male microglia utilize Interferon Regulatory Factor 3 and 7 (IRF3/IRF7) (Fiore et al., 2022). These conclusions parallel increasing evidence of sexual dimorphisms in microglia from development through healthy aging and disease (Guillot-Sestier et al., 2021; Kang et al., 2024; Kerr et al., 2019; Lenz & McCarthy, 2015; Schwarz et al., 2012) and are consistent with data from Alzheimer’s disease and aging rodent models showing female microglia have higher levels of CD68, phagocytosis, and proinflammatory cytokines (Guillot-Sestier et al., 2021; Kang et al., 2024).

TREM2 regulation of phagocytic function resembles known TREM2 actions in Disease Associated Microglia or DAM (Deczkowska et al., 2018; Keren-Shaul et al., 2017; Samant et al., 2024; Wang, 2021), but in contrast to DAM development in other neuropathologies, microglia clustering around dying MNs occurs independently of TREM2. In the facial nucleus, one study suggested that the number of “microglia nodules” is reduced by deletion of tumor necrosis factor (TNF) receptors 1 and 2, and this correlated with MN preservation (Raivich et al., 2002), further confirming these are loci of MN phagocytosis. Despite not being necessary for death cluster formation, TREM2 likely regulates phagocytosis in death cluster microglia by its known binding of many apoptotic signals like externalized phosphatidylserine and nucleic acids (Cannon et al., 2012; Kawabori et al., 2015; Kober et al., 2021; Shirotani et al., 2019; Wang et al., 2015).

Death cluster microglia in the spinal cord may be involve in the removal of a subpopulation of small MNs corresponding to γ-MNs. The fact that survival of postnatal Γ-MNs depends on Glia-Derived Neurotrophic Factor (GDNF) from muscle spindles (Shneider et al., 2009), suggests that Γ-MNs may be especially susceptible to cell death and their loss contributes to the known deficits in proprioceptive mechanisms after nerve injury (Alvarez et al., 2020). However, alternative explanations of our data are possible. The reduced γ-MN cell size peak was accompanied by a larger number of NeuN low-expressing MNs of sizes intermediate between the γ-MN and α-MN peaks. It is thus possible that some γ-MNs have enlarged size after regeneration, or that α-MNs do not fully recover NeuN expression after muscle reinnervation, or that a proportion of MNs reinnervate both muscle spindles and extrafusal muscle fibers forming an expanded β-MN system expressing low NeuN. These interpretative difficulties prevented exact estimations of γ-MN loss and future studies are needed to identify the exact nature of dying MNs using markers that are independent of cell size and NeuN and that withstand changes after axotomy and regeneration. Remarkably changes in the γ-MN peak and the distribution of sizes of MNs with low NeuN were prevented when *trem2* was removed from microglia.

### Conclusions

We characterized two microglia-MN interactions regulated by TREM2. Microglia in “death clusters” surrounding dying/degenerating MNs express greater levels of TREM2 and TREM2 deficiency affects expression of phagocytic markers like CD68. In contrast, TREM2 deficiency disrupts the retrograde MN cell body reaction associated with regenerating and slows muscle reinnervation. These results reveal the complexity of microglia-MN interactions and TREM2 functions that need to be considered when developing TREM2-directed therapeutic interventions to improve recovery following axotomy.

## Supporting information

Supplemental data statistical tables

## Author Contributions

- Designed Research, T.S.P. and F.J.A.
- Performed Research, T.S.P, Z.H-J., E.L.L., D.N.U., V.A.S., and F.J.A.
- Analyzed data, T.S.P., E.L.L., D.N.U., V.A.S., and F.J.A
- Writing – original draft, T.S.P., and F.J.A.
- Writing – review & editing, T.S.P., E.L.L., and F.J.A
- Wrote the paper T.S.P. and F.J.A.

## Conflict of interest statement

The authors disclose that there are no conflicts of interest.

## Acknowledgements

This work was supported by NIH-NINDS grant R01NS111969 (F.J.A.), NIH Training Grant in Integrative Biology T32NS096050-26 (T.S.P.), and Ruth L. Kirschstein NRSA fellowship F31NS130993 (T.S.P.). During this study, P.C.T. was a Summer Opportunity for Academic Research (SOAR) fellow supported by Emory University College. Imaris 3D analyses were performed in the Emory University Integrated Cellular Imaging Core, supported in part by NINDS Core facilities grant P30NS055077. We thank Dr. David Katz (Emory University, Atlanta, GA, USA) for providing the TREM2 KO mouse, Yu Bai (Katz lab) for performing PCR genotyping as needed, and Dr. Bruce Lamb (University of Indiana, Bloomington, IN, USA) for donating the TREM2 flox/flox mouse.

## Abbreviation Key

(CNS): Central Nervous System
(MN): Motoneuron
(PNI): Peripheral Nerve Injury
(CSF1): Colony Stimulating Factor 1
(CSF1-R): CSF1-Receptor
(CCR2): C-C chemokine receptor 2
(TREM2): Triggering Receptor Expressed on Myeloid Cells 2
(DAP12/*tyrobp*): DNAX activating protein of 12kDa
(ePtdSer): externalized Phosphatidylserine
(i.p.): intraperitoneal
(LG): lateral gastrocnemius
(GKO): Global Knockout
(CKO): Conditional Knockout
(WT): Wildtype
(eGFP): enhanced Green Fluorescent Protein
(IHC): Immunohistochemistry
(AAV1-tdTom): Adeno-Associated Virus serotype 1 tdTomato
(P5): postnatal day 5
(dpi): Days Post-Injury
(PFA): paraformaldehyde
(PBS): Phosphate Buffer Solution
(PBST): Phosphate Buffer Solution with 0.03% Triton
(aCSF): artificial Cerebrospinal Fluid
(ROI): Region of Interest
(NMJ): Neuromuscular Junction
(α-BTx): α-Bungarotoxin
(p-SYK): phosphorylated Spleen Tyrosine Kinase
(wpi): Weeks Post-Injury
(DAM): Disease-Associated Microglia
(ChAT): Choline Acetyltransferase
(NeuN): Neuronal Nuclei
(K-S): Kolmogorov-Smirnov
(NFH): Neurofilament Heavy Chain
(VAChT): Vesicular Acetylcholine Transporter
(α-BTx): alpha Bungarotoxin
(Cum-Sum): Cumulative-Summation

## Notes

### Competing Interest Statement

The authors have declared no competing interest.

### Summary of Updates

This revised manuscript reflects the version accepted in the Journal of Neuroscience following review and revisions. JN-RM-0112-26R1 Accepted 27th Mar 2026

