## Supplemental Figures S1 to S7 for "Dual Role of Microglial TREM2 in Neuronal Degeneration and Regeneration After Axotomy"

**Figure S1. Sex effect checks for Tmem119$^{eGFP}$ wildtype (WT) Sholl analysis.**

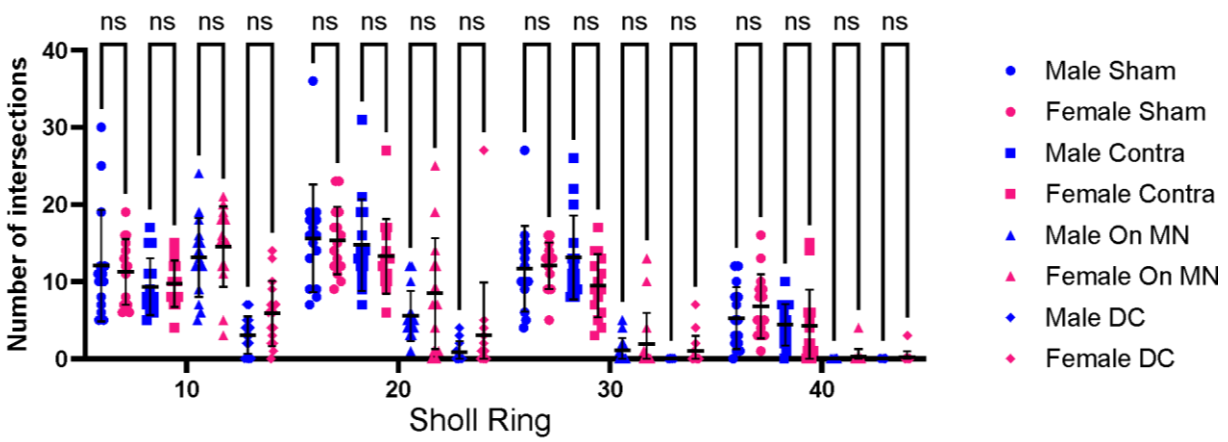

Each dot represents an individual microglia value. Mean ±SD are indicated No sex effect was detected in Sholl analyses. Bonferroni post hoc multiple comparisons, see table below. n= 5 microglia per condition per animal (n=15), N = 3 animals per sex and microglia types.

| Comparison | Mean 1 | Mean 2 | Mean diff. | N1 | N2 | t | DF | P |
|---|---|---|---|---|---|---|---|---|
| Male Sham vs. Female Sham | 12.07 | 11.27 | 0.8 | 15 | 15 | 0.5145 | 448 | >0.9999 |
| Male Contra vs. Female Contra | 9.333 | 9.733 | -0.4 | 15 | 15 | 0.2572 | 448 | >0.9999 |
| Male On MN vs. Female On MN | 13.13 | 14.53 | -1.4 | 15 | 15 | 0.9003 | 448 | >0.9999 |
| Male DC vs. Female DC | 3.067 | 5.867 | -2.8 | 15 | 15 | 1.801 | 448 | >0.9999 |
| Male Sham vs. Female Sham | 15.6 | 15.33 | 0.2667 | 15 | 15 | 0.1715 | 448 | >0.9999 |
| Male Contra vs. Female Contra | 14.73 | 13.27 | 1.467 | 15 | 15 | 0.9432 | 448 | >0.9999 |
| Male On MN vs. Female On MN | 5.533 | 8.467 | -2.933 | 15 | 15 | 1.886 | 448 | 0.9583 |
| Male DC vs. Female DC | 0.8667 | 3.067 | -2.2 | 15 | 15 | 1.415 | 448 | >0.9999 |
| Male Sham vs. Female Sham | 11.67 | 12.07 | -0.4 | 15 | 15 | 0.2572 | 448 | >0.9999 |
| Male Contra vs. Female Contra | 13.13 | 9.467 | 3.667 | 15 | 15 | 2.358 | 448 | 0.3009 |
| Male On MN vs. Female On MN | 1.067 | 1.867 | -0.8 | 15 | 15 | 0.5145 | 448 | >0.9999 |
| Male DC vs. Female DC | 0 | 1 | -1 | 15 | 15 | 0.6431 | 448 | >0.9999 |
| Male Sham vs.Female Sham | 5.267 | 6.8 | -1.533 | 15 | 15 | 0.9861 | 448 | >0.9999 |
| Male Contra vs. Female Contra | 4.4 | 4.267 | 0.1333 | 15 | 15 | 0.08574 | 448 | >0.9999 |
| Male On MN vs. 40:Female On MN | 0 | 0.2667 | -0.2667 | 15 | 15 | 0.1715 | 448 | >0.9999 |
| Male DC vs. Female DC | 0 | 0.2 | -0.2 | 15 | 15 | 0.1286 | 448 | >0.9999 |

**Figure S2. Sex effect checks TREM2 expression (IHC) in Tmem119$^{eGFP}$ microglia.**

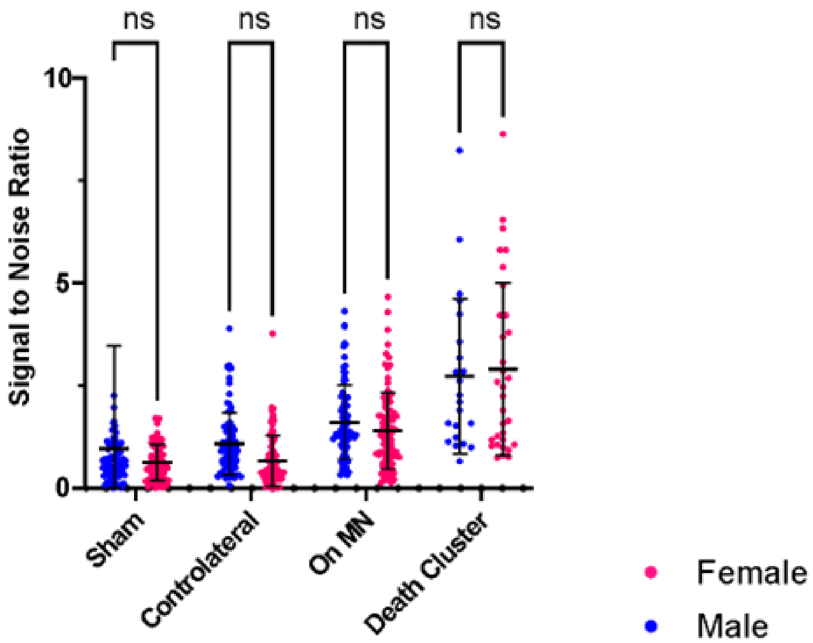

No sex effect was detected in TREM2 immunofluorescence (signal to noise ratio) on the membrane and cytosol of different eGFP labeled microglia. Values are individual microglia (n=17-26 microglia per animal per condition, N= 4 per sex per morphology) and analyzed via Bonferroni post hoc multiple comparisons.

| **Comparison** | **mean 1** | **mean 2** | **mean diff.** | **N1** | **N2** | **t** | **DF** | **P** |
|---|---|---|---|---|---|---|---|---|
| Male - Female | | | | | | | | |
| Sham | 0.9642 | 0.6196 | 0.3446 | 81 | 80 | 1.651 | 549 | 0.3974 |
| Contralateral | 1.088 | 0.6644 | 0.4232 | 84 | 78 | 2.033 | 549 | 0.1703 |
| On MN | 1.611 | 1.399 | 0.2111 | 85 | 95 | 1.068 | 549 | >0.9999 |
| Death Cluster | 2.73 | 2.901 | -0.1713 | 22 | 32 | 0.467 | 549 | >0.9999 |

**Figure S3. Sex effect checks *trem2* mRNA expression (RNAscope) in CX3CR1$^{eGFP}$ microglia.**

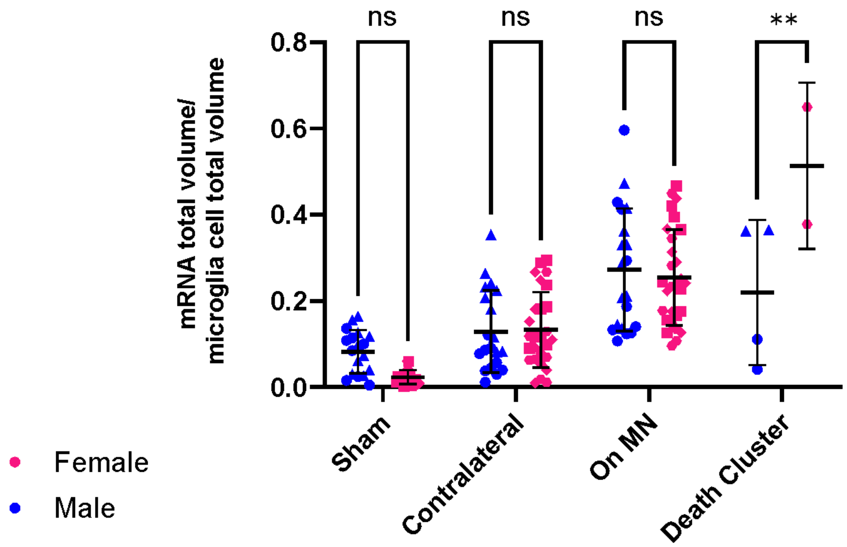

*trem2* mRNA signal in 3D reconstructed individual eGFP-labeled microglia (n=10 microglia per animal per condition, N=2 sham per sex, 5 cut-ligation (2 males, 3 females). Mean ±SD of volume occupancy is shown. Shown are results from Dunn's post hoc multiple comparisons on n = number of microglia (**p=0.0029) (see table below).

| **Comparison** | **mean 1** | **mean 2** | **mean diff.** | **N1** | **N2** | **t** | **DF** | **P** |
|---|---|---|---|---|---|---|---|---|
| Male - Female | | | | | | | | |
| Sham | 0.0823 | 0.02307 | 0.05923 | 18 | 20 | 1.858 | 132 | 0.2619 |
| Contralateral | 0.1289 | 0.1328 | -0.003854 | 20 | 27 | 0.1331 | 132 | >0.9999 |
| On MN | 0.2723 | 0.2545 | 0.0178 | 20 | 29 | 0.6239 | 132 | >0.9999 |
| Death Cluster | 0.2198 | 0.5138 | -0.294 | 4 | 2 | 3.459 | 132 | 0.0029 |

**Figure S4. Sex checks on phospho-SYK (p-SYK) expression in different types of microglia in Tmem119$^{eGFP}$ wildtype (WT) and TREM2 Conditional Knock Out (CKO) mice.**

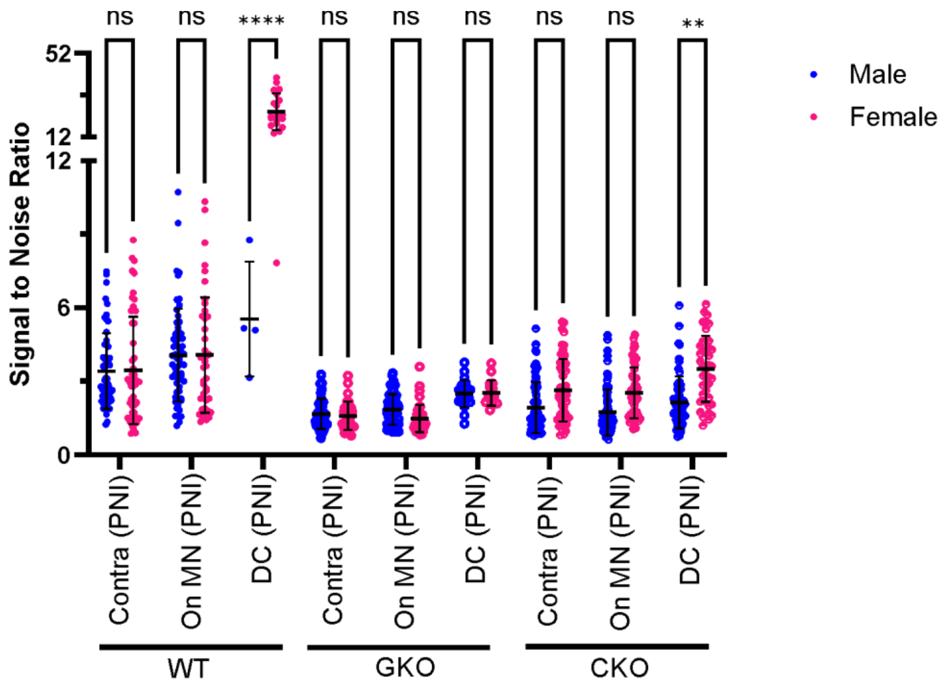

Signal-to-noise ratio of microglia soma membrane p-SYK immunofluorescence in different types of microglia (contralateral, On MN and DC). Each dot represents one microglia and mean ±SD are represented; n=17-20 microglia per condition per animal, N= 6 WT cut-ligation (equal sex), 5 GKO cut-ligation (3 males, 2 females), 7 CKO cut-ligation (4 males, 3 females). Dunn's post hoc comparisons based on n, number of microglia analyzed (**p=0.0022, ****p<0.0001).

| Comparison | mean 1 | mean 2 | mean diff. | N1 | N2 | t | DF | P |
|---|---|---|---|---|---|---|---|---|
| Male - Female | | | | | | | | |
| WT Contralateral (PNI) | 3.413 | 3.445 | -0.03164 | 58 | 50 | 0.08776 | 803 | >0.9999 |
| WT On MN (PNI) | 4.062 | 4.073 | -0.0103 | 61 | 43 | 0.02768 | 803 | >0.9999 |
| WT Death Cluster (PNI) | 5.539 | 24.05 | -18.51 | 4 | 20 | 18.08 | 803 | <0.0001 |
| GKO Contralateral (PNI) | 1.668 | 1.584 | 0.08365 | 53 | 33 | 0.2019 | 803 | >0.9999 |
| GKO On MN (PNI) | 1.851 | 1.482 | 0.369 | 59 | 40 | 0.9642 | 803 | >0.9999 |
| GKO Death Cluster (PNI) | 2.493 | 2.524 | -0.03083 | 22 | 16 | 0.05022 | 803 | >0.9999 |
| CKO Contralateral (PNI) | 1.926 | 2.629 | -0.7027 | 71 | 55 | 2.094 | 803 | 0.3294 |
| CKO On MN (PNI) | 1.731 | 2.529 | -0.7976 | 75 | 55 | 2.405 | 803 | 0.1478 |
| CKO Death Cluster (PNI) | 2.143 | 3.511 | -1.368 | 64 | 42 | 3.688 | 803 | 0.0022 |

**Figure S5. Sex effects for microglia number and territory size in uninjured animals that are either wildtype for *trem2* (WT, Tmem119$^{eGFP}$ microglia), global *trem2* KO (GKO, Tmem119$^{eGFP}$ microglia) or conditional t*rem2* KO (CKO.,Tmem119$^{CreERT2}$:: tdTomato microglia).**

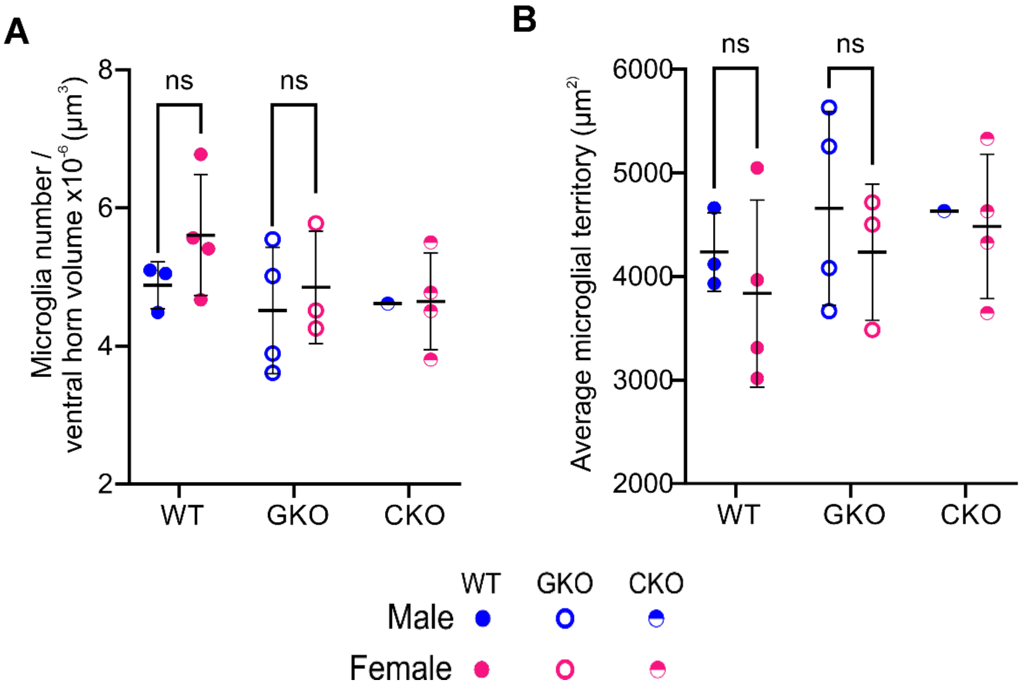

**A)** Number of ventral horn microglia in 100 µm side cubes estimated in five L5 sections per animal. Each dot represents the animal average, mean ±SD represented. No statistical difference between sexes (Bonferroni post-hoc tests, see table below). **B)** Similarly, no differences were found on average territory area. Same sample and statistical comparison as in A. Note no statistical comparison performed for CKO because only one male was analyzed.

| **Comparison** | **mean 1** | **mean 2** | **mean diff.** | **N1** | **N2** | **t** | **DF** | **P** |
|---|---|---|---|---|---|---|---|---|
| Male-Female | | | | | | | | |
| **Microglia Number (A)** | | | | | | | | |
| WT | 4.878 | 5.606 | -0.7276 | 3 | 4 | 1.227 | 13 | 0.7246 |
| GKO | 4.515 | 4.849 | -0.3344 | 4 | 3 | 0.564 | 13 | >0.9999 |
| CKO (N/A) | 4.615 | 4.646 | -0.03164 | 1 | 4 | N/A | N/A | N/A |
| **Microglia territories (B)** | | | | | | | | |
| WT | 4237 | 3835 | 402.5 | 3 | 4 | 0.686 | 13 | 0.6758 |
| GKO | 4659 | 4233 | 425.4 | 4 | 3 | 0.7252 | 13 | 0.6758 |
| CKO (N/A) | 4634 | 4483 | 150.4 | 1 | 4 | N/A | N/A | N/A |

**Figure S6. Sex effects for the number of death clusters in injured animals that are trem2 wildtypes (WT, Tmem119$^{eGFP}$ microglia), global *trem2* KOs (GKO, Tmem119$^{eGFP}$ microglia) or conditional t*rem2* KOs (CKO.,Tmem119$^{CreERT2}$:: tdTomato microglia).**

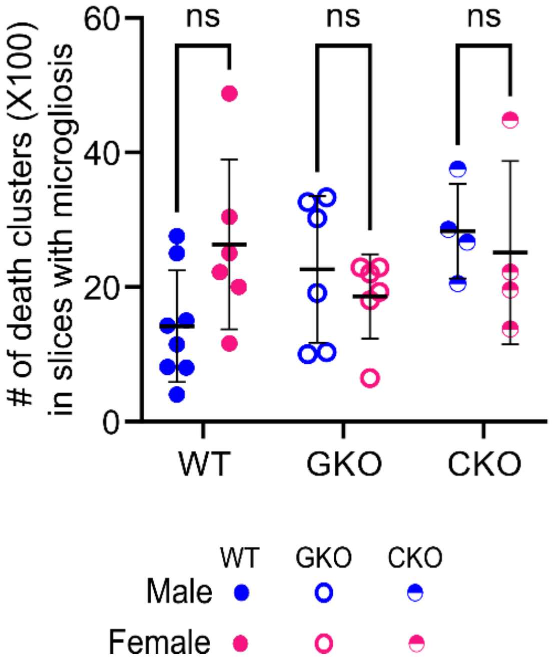

No sex differences were detected in the number of death cluster detected in males and females. The number of death clusters throughout an animal's entire lumbar spinal cord was divided by the number of sections with apparent microgliosis for standardization and multiplied by 100 to acquire the total number on 5 mm column. Each dot is one animal estimate. The animals mean ±SD are represented and analyzed with Bonferroni post hoc multiple comparisons.

| Comparison | mean 1 | mean 2 | mean diff. | N1 | N2 | t | DF | P |
|---|---|---|---|---|---|---|---|---|
| Male-Female | | | | | | | | |
| WT | 14.18 | 26.34 | -12.16 | 8 | 6 | 2.262 | 28 | 0.0949 |
| GKO | 22.61 | 18.6 | 4.009 | 6 | 6 | 0.6974 | 28 | >0.9999 |
| CKO (N/A) | 28.31 | 25.1 | 3.211 | 4 | 4 | 0.4561 | 28 | >0.9999 |

**Figure S7. Sex effects for the number of microglia-MN interactions in injured animals that are trem2 wildtypes (WT, Tmem119$^{eGFP}$ microglia), global *trem2* KOs (GKO, Tmem119$^{eGFP}$ microglia) or conditional t*rem2* KOs (CKO.,Tmem119$^{CreERT2}$:: tdTomato microglia).**

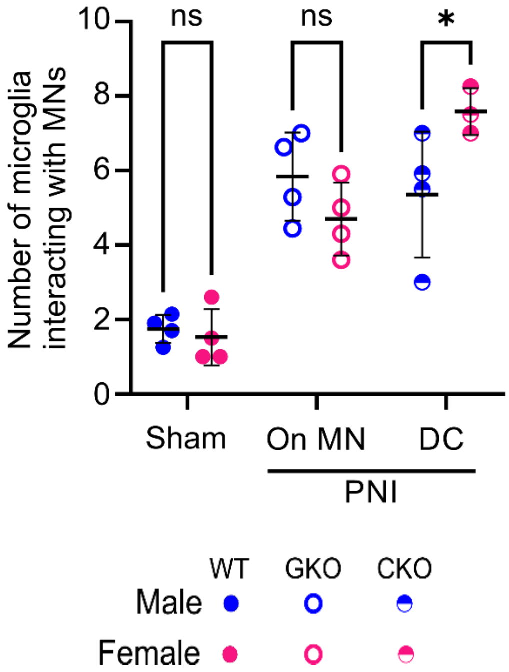

Females had significantly larger number of microglia forming part of death cluster in Tmem119 eGFP wildtype mice (*p=0.037, Bonferroni post-hoc-test). There were no sex differences in the number of microglia-MN contacts in sham animals or microglia On MN while they regenerate (n= 7-10 MNs per animal per condition, N=4 animals per condition). Each dot is one animal estimate. The animals mean ±SD are represented and analyzed with Bonferroni post hoc multiple comparisons.

| Comparison | mean 1 | mean 2 | mean diff. | N1 | N2 | t | DF | P |
|---|---|---|---|---|---|---|---|---|
| Male-Female | | | | | | | | |
| WT | 1.748 | 1.525 | 0.2232 | 4 | 4 | 0.3015 | 17 | >0.9999 |
| GKO | 5.839 | 4.7 | 1.139 | 4 | 4 | 1.538 | 17 | 0.4272 |
| CKO (N/A) | 5.354 | 7.583 | -2.229 | 4 | 3 | 2.788 | 17 | 0.0379 |
